# High-resolution microCT reveals relationships between stomata and interior leaf anatomy in Sorghum

**DOI:** 10.64898/2025.12.05.692612

**Authors:** James M. Fischer, Guillaume Théroux-Rancourt, Craig R. Brodersen, Andrew D.B. Leakey

## Abstract

Stomata are pores in the leaf epidermis that regulate the trade-off between CO_2_ uptake for photosynthesis and water vapor loss to the atmosphere. Stomatal patterning therefore influences water use efficiency and is a target for engineering to avoid drought stress. However, there is limited understanding of how internal leaf anatomy is coordinated with stomatal development, in part due to the technical challenges of assessing three-dimensional anatomy with sufficient resolution. C4 grasses are understudied, and this is a significant knowledge gap given their file-like stomatal distribution and unique mesophyll organization. In this study, wild-type sorghum and a low-stomatal density transgenic line expressing a synthetic Epidermal Patterning Factor (EPF_syn_) were studied. High-resolution microCT was paired with machine learning to characterize three-dimensional traits of mesophyll, epidermis, and airspace, which together determine g_ias_. Sorghum internal leaf airspace is an arrangement of large sub-stomatal airspaces with thin air passageways. Adaxial and abaxial surfaces differed in stomatal patterning relative to mesophyll structures, sub-stomatal crypts and airspace CO_2_ conductance (g_ias_). Surprisingly, adaxial stomata were consistently located above rather than between vascular bundles. Unexpectedly, g_ias_ was not significantly different in wild-type versus EPF_syn_. EPF_syn_ plants had larger crypts and shifts in internal leaf anatomy, indicating a potential compensation mechanism for predicted impacts of reduced stomatal density on g_ias_. These findings provide a new understanding of the interplay between leaf surface specific anatomy and internal structural patterning of the mesophyll in a C4 species, and provides knowledge relevant to engineering water use efficiency in crop species.

## INTRODUCTION

Crop water use efficiency (WUE) at the whole-plant scale is defined as biomass production per unit of water use, and has been a key crop improvement target for over a century because water is the most important environmental factor limiting crop production (Boyer, 1982; Briggs and Shantz, 1917; Condon et al., 2004; Gray and Dunn, 2024; Leakey et al., 2019). Recently, significant advances have been made in understanding how intrinsic WUE (iWUE; photosynthetic CO_2_ uptake (*A*) relative to stomatal conductance (*g_s_*)) at the leaf scale can be improved by engineering reductions in stomatal density (SD) or altering stomatal behavior (Caine et al., 2019; Ferguson et al., 2024; Franks et al., 2015; Hughes et al., 2017; Lunn et al., 2024; Schuler et al., 2018; Shahbaz et al., 2025). The underlying rationale for such efforts is that iWUE can be improved by reducing water loss with little or no loss in carbon gain. This is particularly true in C4 species, which possess a carbon-concentration mechanism (CCM) (von Caemmerer and Furbank, 2003; Ferguson et al., 2024). Without a CCM, reductions in SD would reduce water loss, but at the expense of sub-saturating [CO_2_] inside the leaf because of the reduction in the potential entry points for CO_2_ diffusion across the epidermis (von Caemmerer and Furbank, 2003; Ferguson et al., 2024). Efforts to improve WUE—by manipulating stomatal density and patterning—leverage understanding of the developmental coordination of epidermal cells and the underlying mesophyll, vasculature, bundle sheath, veins, and airspaces (Baillie and Fleming, 2019; Lundgren et al., 2019; Pathare et al., 2020; Pathare et al., 2024; Trueba et al., 2022; Whitewoods, 2021). However, significant knowledge gaps remain about the interactions between stomatal patterning and the three-dimensional anatomy of leaves, especially in grasses and species with C4 photosynthesis.

Entering through the open stomate from the external atmosphere, CO_2_ then diffuses through the interior airspace to the sites of CO_2_ fixation in the mesophyll. During leaf development, proto-mesophyll cells located below immature stomates divide and sculpt, separating from one another to form as the airspaces of the “sub-stomatal crypts” (Fig. 1A, B; Baillie and Fleming, 2019). These crypts create the bulk of porous space within the leaf and are primary diffusive pathways for CO_2_ (de Oliviera et al., 2023). However, the presence of stomata embedded in the epidermis and the corresponding sub-stomatal crypts necessarily displace other tissue types, such as photosynthetic mesophyll cells, veins, or epidermal structures (e.g. large bulliform cells) (Baresch et al. 2018; Karabourniotis et al., 2021; Noblin et al., 2008). A major outstanding question is whether there is a structural optimum which balances the partitioning of the leaf volume to meet the multiple demands imposed on the leaf that satisfy its photosynthetic physiology, water transport, photosynthate export, and biomechanical needs, all while maximizing CO_2_ uptake and minimizing water loss (Drake et al., 2018; Earles et al., 2018; Noblin et al., 2008; de Oliveira et al., 2022). As such, coordination between stomata and the leaf interior—beyond the subtending air crypts—has been shown to guide patterning of the leaf (Baillie and Fleming, 2019; Lundgren et al., 2019; Pathare et al. 2020; Pathare et al. 2024; Trueba et al. 2022; Whitewoods, 2021). For example, stomatal files are thought to avoid forming over vasculature, where the development of associated internal airspace would be impeded by the limited space between epidermis and underlying bundle sheath (McKown and Bergmann, 2020; Schuler et al., 2018). Understanding relationships between internal leaf components (e.g. airspace) and external leaf components (e.g. stomata) remain important in understanding leaf physiology. A major limitation in quantifying and studying the relationships between leaf structure and function has been the capacity for producing accurate, three-dimensional datasets with sufficient resolution to accurately characterize the absorptive surfaces and diffusive pathways inside the leaf.

**Fig. 1:**
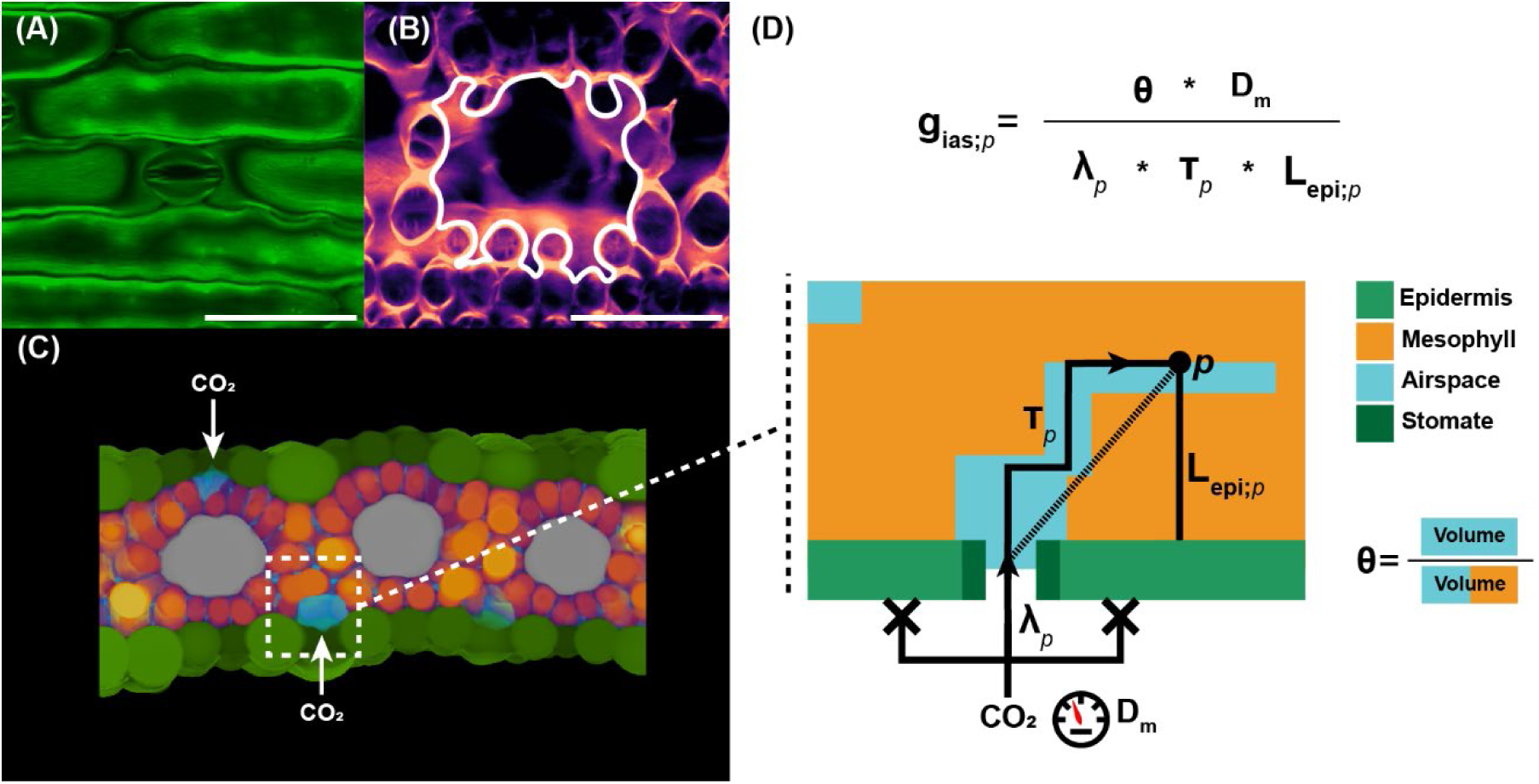
(A) Confocal scan of *Sorghum bicolor* abaxial stomatal complex (B) mesophyll and airspace underlying stomate in A, white outline indicates boundaries of sub-stomatal air crypt (C) cross-sectional cartoon of sorghum leaf, with arrows indicating stomata and sub-stomatal crypts (D) simplified cartoon and calculations of airspace conductance to CO_2_ at point *p* at mesophyll/airspace contact. See Materials and Methods for formulas for calculating g_ias_ and components.

Recently, high-resolution X-ray micro-computed tomography (microCT) has been developed as a means for investigating the three-dimensional organization of plant structures without compromising the architectural integrity of the tissue (Earles et al., 2018; Harwood et al., 2021; Mathers et al., 2018; Piovesan et al., 2021; Théroux-Rancourt et al., 2021). The three-dimensional approach of microCT has created new opportunities to study leaf anatomy and structure-function relationships at sub-micron resolutions, such as quantifying inherently three-dimensional traits which influence internal CO_2_ movement (Borsuk et al. 2022; Borsuk et al., 2025; Earles et al., 2018; Earles et al., 2019; Théroux-Rancourt et al., 2023). For example, microCT has enabled the discovery of gradients of porosity (θ)—a ratio relating airspace volume and mesophyll volume—across the leaf depth, with greater θ often occurring in the abaxial region of the leaf (Baillie and Fleming, 2019; Dorca-Fornell et al., 2013; Lehmeier et al., 2017; Mathers et al., 2018). High θ has been associated with higher values of CO_2_ conductance, however other parameters—such as the mesophyll surface area exposed to the IAS—are likely equally or more important, but these parameters are difficult to calculate with two-dimensional data (Earles et al., 2018; Lehmeier et al., 2017; Lundgren et al., 2019; Théroux-Rancourt et al., 2021). Thus, investigating C4 internal leaf anatomy and g_ias_ at a detailed three-dimensional level is crucial to understanding overall mesophyll conductance—of which g_ias_ is a component—and enables more precise engineering of photosynthesis (Earles et al., 2019; Gago et al., 2020; Lehmeier et al., 2017; Trueba et al., 2021).

Prior to the availability of high-resolution 3D data, calculations of internal airspace conductance to CO_2_ (g_ias_) were limited to using challenging techniques such as helox (Mott and Parkhurst 1990; Piel 2002) or resolving to use one and two-dimensional anatomical data on traits that could be estimated with limited dimensionality and image resolution—such as θ—to estimate conductance (Morison and Lawson, 2007). More recent anatomical models of g_ias_, utilizing three-dimensional data, also consider the tortuosity and lateral path lengthening of the airspace to more accurately calculate g_ias_ (Earles et al., 2018; Harwood et al., 2021; Fig. 1D). Lateral path lengthening (λ) accounts for the requirement of CO_2_ entry into the leaf through stomates, and not through the entirety of the epidermis as was assumed by early models of g_ias_ (Earles et al., 2018). If there are no compensatory changes in internal structure, more sparsely distributed stomata (lower SD) would increase the average distance from the stomates to the diffusive surface of mesophyll cells, hence increasing λ and decreasing g_ias_.

Tortuosity (τ)—a second component of g_ias_—accounts for the complexity of movement through the airspace from the entry point of a stomate (e.g. movement around bulbous cells blocking direct paths further into the airspace). Harwood et al. (2021) found that increased SD can drive lower τ by increasing the number of sub-stomatal air crypts, thereby reducing the path length from stomate to the diffusive surface. Lundgren et al. (2019) found that θ was significantly lower in a reduced-SD wheat line, which would theoretically act to lower g_ias_. A reduction in θ concomitant with SD-reduction is logical, as decreasing the number of stomata inherently decreases the number of sub-stomatal crypts, a major driver in leaf porosity (Hughes et al., 2017). This would add resistance to CO_2_ diffusion from the stomata through the internal airspace to mesophyll cells, and would decrease CO_2_ concentrations at the site of CO_2_ fixation. Similarly, SD changes also drive shifts in mesophyll cell density and patterning, which can then in turn affect the τ of the airspace (e.g. smaller, more densely packed mesophyll cells could increase difficulty of movement through the airspace) (Dow et al., 2017). However, λ, τ, and g_ias_ were not estimated in prior studies of low-SD plants, so the prediction of lower g_ias_ associated with lowered SD is tentative and needs further investigation.

To achieve a more holistic understanding of relationships between epidermal stomatal patterning, internal leaf anatomy, and CO_2_ conductance we designed this study to investigate how changes in SD influence the organization of the underlying mesophyll tissue and airspace, and how those structural changes might alter airspace conductance. We utilized a transgenic line of sorghum (*Sorghum bicolor*) expressing a synthetic Epidermal Patterning Factor (EPF_syn_; Ferguson et al., 2024). In previous work, EPF_syn_ plants averaged a 45-69% reduction in SD, approximately 30% reduction in water use, and a 13-20% increase in iWUE (Ferguson et al., 2024). Working within this framework we tested the hypotheses that: **(1) internal airspace on the abaxial versus adaxial sides of the leaf will consistently be concentrated between vascular bundles; (2) adaxial and abaxial g_ias_ will vary in proportion to the stomatal density differences between sides; (3) plants with low-SD driven by expressing EPF_syn_ will have lower θ and g_ias_ than wild-type.** High-resolution microCT images were analyzed with a newly trained machine learning pipeline specific to sorghum, which enabled construction of a three-dimensional leaf structure with sub-micrometer resolution. Sorghum is a model C4 species and an important crop for production of food, forage, and biomass, which can be converted into biofuel and bioproducts (Castro et al., 2017; Morris et al., 2013). As a C4 species, sorghum makes an ideal species for answering these questions because reductions in SD achieved through EPF_syn_ expression should only limit water loss, while the C4 CCM maintains CO_2_ at sufficiently high concentrations to support photosynthesis (von Caemmerer and Furbank, 2003; Ferguson et al., 2024).

## METHODS

### Plant growth

Wildtype (WT) grain *Sorghum bicolor* genotype Tx430 and a transgenic line ubiquitously expressing a synthetic EPF (EPF_syn_) which has been previously described (Ferguson et al., 2024), were grown under greenhouse conditions until 4-5 fully expanded leaves were present. Six to eight replicate plants of each genotype were grown in a randomized block design. Daytime temperature was 28-30°C and nighttime temperature was 21-23°C. Supplemental lighting maintained a day length of 15 hours and a minimum daytime light irradiance of 1,290 µmol s^-1^ m^-2^.

### X-ray imaging, reconstruction, and machine learning labeling

MicroCT scanning was performed at the Advanced Photon Source (APS) beamline BM-2 at Argonne National Laboratory (Lemont, IL, USA). Leaf tissue was harvested fresh from the plant, approximately 15 minutes prior to scanning. Samples were harvested from the newest fully-expanded leaf (leaf 4-5), at a central location halfway between base and tip. Two to three sub-samples, extending from the mid-rib to leaf margin at a midway point along the leaf, were harvested per leaf. Samples were individually wrapped in polyimide tape to prevent drying (Earles et al., 2018). Tape-wrapped samples were placed into a clamp and scanned, with each 180-degree rotational scan taking ∼15 minutes, collecting ∼1500 scans per sample. A 10x objective was used with a resolution of 0.65 µm/pixel.

Projections obtained from the beamline were reconstructed in the cross-sectional plane using APS’ in-house reconstruction platform TomoPy (Gürsoy et al., 2014). For enhanced contrast, phase and non-phase channels were reconstructed then merged in ImageJ. Reconstructed scans were then rotated so that the leaf surfaces were parallel with the x-axis in the cross-sectional plane.

For machine-learning based labeling of tissues, 20 raw slices were selected from the pool of samples, and hand labeled in ImageJ. Four different classes were labeled: epidermis, airspace, mesophyll, and vascular bundle (including both veins and bundle sheath cells), with each class assigned a different fill color. Hand-labeled slices were used to train a mask-RCNN machine learning (ML) pipeline in PyTorch (Paszke et al., 2019). The trained mask-RCNN model was used to automatically label all slices within each image stack. ML-labeled stacks were trimmed to remove damaged tissues. After trimming, samples (in the cross-sectional plane) had an average width of 1,009 µm and average length of 1,426 µm, for an average area of 1,429,625 µm^2^ (1.43 mm^2^).

### Manual stomatal density and stomate dimension measurements

Stomates were manually labeled in ML-stacks by placing a dot over the stomatal pore in ImageJ and recording x, y, and z coordinates. Stomate locations and leaf area were used to manually calculate stomatal density (SD). Stomate coordinates were used in downstream analyses, described below.

For each ML-labeled stack, a blank stack with the same dimensions was created in ImageJ, and resliced to the paradermal plane. Stomates were labeled with a dot placed in the center of the stomate at the point where airspace and stomatal pore meet in the paradermal plane.

Individual stomate dimensions (length, width, area) were calculated in ImageJ/FIJI by manually drawing ovals around individual stomates in the paradermal plane, then these ovals were measured with FIJI’s Particle Analyzer (Schindelin et al., 2012). Stomates were labeled adaxial or abaxial based on centroid location. Stomatal files (and therefore stomatal length) ran parallel to the x-axis of the paradermal stack.

### Stomate and vascular bundle “contact” analysis

Stomatal coordinates (x, y, z) were extracted from the tagged stomatal locations in ImageJ/FIJI. Stomates were grouped into adaxial and abaxial classes based on their coordinates. Stomatal coordinates and vasculature binary mask stacks were fed into R (R Core Team, 2021). A straight vertical line for each stomate was created using the stomatal coordinates and stack dimensions, and R packages sf, sp, and rgeos (Pebesma and Bivand, 2005; Pebesma and Bivand, 2023). Vertical lines were merged with binary vascular masks, and the number of stomatal lines that intersected vascular bundles were counted per sample.

To split ML-labeled masks into adaxial and abaxial regions, the centroids and bounding boxes of vascular bundles were determined on a per-slice basis in the cross-sectional plane, using Image’s Particle Analyzer (Schindelin et al., 2012). A horizontal line was drawn in the cross-sectional plane, connecting the centroids of the vascular bundles in the lateral direction. Regions above this horizontal line were labeled adaxial, and regions below the line were labeled abaxial.

### Volumes, mesophyll surface area, and airspace connectivity

Three-dimensional volumes and surface areas of the four different ML-labeled classes (epidermis, vascular bundles including veins and bundle sheath cells, airspace, mesophyll) were measured with BoneJ Particle Analyzer (Doube et al., 2010). See Table 1 for keywords and definitions summarized from the following sections. Porosity (θ) was calculated as the volume of airspace as a ratio to the sum of airspace and mesophyll volumes (V_m_ = V_ias_ + V_meso_):

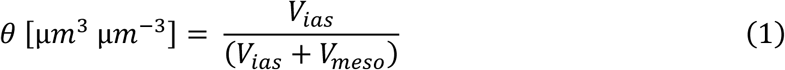

**Table 1:** Key terms related to 3D leaf airspace morphology and physiology. See Materials and Methods for more detailed descriptions.

| ***term*** | ***unit*** | ***definition*** |
| --- | --- | --- |
| **Contact surface area ($S_m$)** | $\mu\text{m}^2$ | The surface area of the region where the mesophyll contacts the intercellular airspace |
| **Connectivity ($K_{SA}$)** | $\mu\text{m}^3 \mu\text{m}^{-3}$ | Fraction of the largest air volume connected to the external atmosphere |
| **$V_m$** | $\mu\text{m}^3$ | Sum of intercellular airspace volume ($V_{ias}$) and mesophyll volume ($V_{mes}$) |
| **$S_{m,LA}$** | $\mu\text{m}^2 \mu\text{m}^{-2}$ | Ratio between $S_m$ and leaf area (LA) |
| **$S_{m,V}$** | $\mu\text{m}^2 \mu\text{m}^{-3}$ | Ratio between $S_m$ and $V_m$ |
| **Porosity ($\theta$)** | $\mu\text{m}^3 \mu\text{m}^{-3}$ | Ratio between airspace volume ($V_{ias}$) and $V_m$ (Formula 1) |
| **$L_{mes}$** | $\mu\text{m}$ | Vertical height of the mesophyll from adaxial to abaxial surfaces |
| **$L_{epi}$** | $\mu\text{m}$ | Distance from epidermises to any point in the airspace, NOT considering obstacles |
| **$L_{euc}$** | $\mu\text{m}$ | Distance from stomate to any point in the airspace, NOT considering obstacles |
| **$L_{geo}$ (Diffusive path length)** | $\mu\text{m}$ | Distance from stomate to any point in the airspace, considering obstacles (e.g. mesophyll) |
| **Lateral path lengthening ($\lambda$)** | $\mu\text{m} \mu\text{m}^{-1}$ | Difficulty in lateral movement through the airspace (Formula 5) |
| **Tortuosity ($\tau$)** | $\mu\text{m} \mu\text{m}^{-1}$ | Difficulty of movement from stomate to mesophyll surface through the airspace (Formula 4) |
| **$g_{ias}$** | $\text{mol m}^{-2} \text{s}^{-1}$ | Intercellular airspace conductance of $\text{CO}_2$ (Formulas 6-7) |

Leaf area (A_leaf_), V_m_, and total mesophyll surface area contacting the airspace (S_m_) were used to calculate S_m,V_ and S_m,LA_ using the following equations (Earles et al., 2018; Théroux-Rancourt et al., 2023):

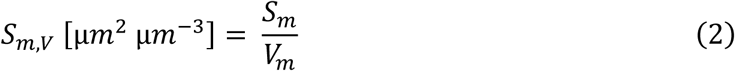

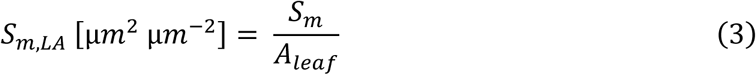

Connectivity (κ_V_) was calculated as the ratio between the volume of the largest airspace particle in the sample, and the sum of total airspace volume, converted to a percentage (Earles et al., 2018). Surface area was calculated for each air particle using BoneJ’s Particle Analyzer (Doube et al., 2010). As major veins completely bisected the airspace into two distinct particles (Supp. fig. 1), only samples with no major veins in the image were used to calculate κ_V_ to avoid bias.

Pathway analyses (Supp. fig. 4) were performed with FIJI/Flood Tools, first calculating the geodesic distance between a set of points (e.g. adaxial stomates) and the airspace, then using the geodesic distance map and a second set of points (e.g. abaxial stomates) to calculate the shortest paths between the two sets of points through the airspace. These paths were analyzed with FIJI/BoneJ tool Analyze Skeleton, and geodesic/Euclidian distances were used to calculate tortuosity of the path (equation 4). The binary paths were merged with the airspace diameter stack for the sample to calculate the diameters of the air channels that the paths passed through from one point to the other.

### Crypt isolation

To extract sub-stomatal air crypts, ML-labeled stacks were resliced to the paradermal plane, with vasculature running parallel to the x-axis of the stack. The average diameter of the full airspace for each sample was calculated with FIJI’s local thickness plugin, as the average diameter of all slices within the stack (Diam_full air_). To remove thin interconnecting airspaces between crypts a tophat filter was applied with the threshold equal to Diam_full air_ * 1.75. Factor 1.75 was selected as this value proved to isolate crypts most accurately compared to other tested values; this value is likely species and experiment dependent. As genotypes did not differ in overall airspace diameter, airspace diameter was used to determine the threshold cutoff for crypt isolation. A marker-controlled watershed was then run via FIJI/MorpholibJ, with stomatal points as seeds on the tophat filtered airspace to extract air crypts directly associated with stomates (Supp. fig. 3; Legland et al., 2016).

Extracted air crypts were analyzed with FIJI/3dSuite’s 3D Manager (Ollion et al., 2013) for volume and bounding box dimensions. Length, width, and depth were calculated from the bounding box dimensions of each crypt in the x, y, and z dimensions respectively. Crypts were labeled adaxial or abaxial based on centroid location. Crypts were subtracted from the original airspace mask to create a non-crypt airspace stack.

### Airspace conductance (g_ias_)

The following analyses to derive conductance traits of interest follow well-established protocols for analyzing high-resolution X-ray data of leaf anatomy (Earles et al., 2018; Harwood et al., 2021; Mathers et al., 2018; Théroux-Rancourt et al., 2021). All code needed to recapitulate analyses will be available upon acceptance of the manuscript at [INSERT WEBSITE URL].

To quantify the contact between internal airspace and mesophyll, the binary mesophyll mask was dilated one pixel, and the overlap between mesophyll and airspace was created using FIJI Image Calculator. This mask of mesophyll/airspace contact was used to limit the following calculations only to the mesophyll surface contacting intercellular airspace.

Diffusive path length (L_geo_), tortuosity, and lateral path lengthening were calculated following the approach of Earles et al. (2018). L_geo_ was calculated as the geodesic distance from a stomate to any point along the mesophyll, considering obstacles (e.g. other mesophyll cells), calculated with morpholibJ’s 3D geodesic distance plugin (Legland et al., 2016). Similarly, the unobstructed Euclidean distance (i.e. straight path length; L_euc_) from stomate to mesophyll surface was calculated. The tortuosity factor (τ) of the airspace was calculated per pixel as the squared ratio between the obstructed (L_geo_) and unobstructed distances from stomate to the mesophyll surface (L_euc_):

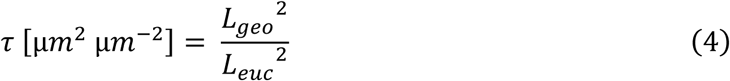

The distance from the surface of the mesophyll to the closest epidermis (L_epi_) was calculated with morpholibJ’s 3D geodesic distance (Legland et al., 2016). Lateral path lengthening (λ) was calculated as the ratio between the Euclidean distance from stomate to mesophyll surface (L_euc_) and L_epi_, to describe the limitations of stomates to gas exchange:

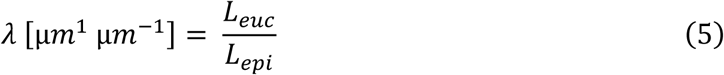

The analyses described above resulted in 3D calculated maps of L_geo_, τ, and λ for each sample. These maps were then divided into adaxial and abaxial regions using vascular centroids. Whole sample, adaxial, and abaxial averages were calculated for τ and λ, and in addition to θ, were used to calculate g_ias_ (m s^-1^):

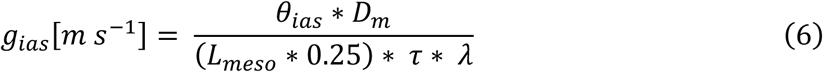

in which L_meso_ is the average cross-sectional thickness of the mesophyll (i.e. leaf thickness without epidermises) multiplied by 0.25 to account for amphistomaty (Harwood et al. 2021). D_m_ is the diffusivity of CO_2_ in air (1.51E^-5^ m^2^ s^-1^). To convert g_ias_ (m s^-1^) to molarity, the equation from Niinements and Reichstein (2003) was used:

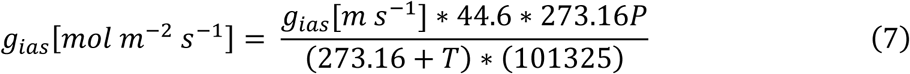

in which P is air pressure (101,325 Pa) and T is air temperature (25°C). Before calculating full sample values, stacks were cropped on the edges of the sample to remove regions near the edges of the stack, to reduce edge bias. Stacks were cropped to remove 1/8^th^ of the stack on each edge of the sample. Stacks were then average z-projected in FIJI, and the mean value of the z-projection calculated as the sample mean for that trait.

### Leaf cross-sectional profiles of airspace and conductance traits

To measure airspace (full airspace and non-crypt airspace) and mesophyll diameters throughout the depth of the leaf, binary masks were resliced to the paradermal view. After reslicing, the masks were analyzed with FIJI/BoneJ’s Slice Geometry tool (Doube et al., 2010). Slices were then scaled from 1 to 100, with 1 being the first slice in which airspace appears on the adaxial side and 100 being the last slice in which airspace appears on the abaxial side.

To express conductance traits across leaf depth, trait stacks created using methods described above were resliced to the paradermal plane, then the FIJI “Plot Z-axis Profile” tool was used to obtain per-slice averages of the conductance traits.

The stat_smooth function from ggplot2 was used to generate average trait profiles across the leaf (Wickham, 2016). First, technical replicates were averaged into leaf averages, i.e. for each biological replicate across the cross-sectional profile. Second, replicate leaves were averaged into treatment means, again across the cross-sectional profile.

To calculate g_ias_ on a per-slice basis, equation 6 was used with per-slice average values for all anatomical traits input so that:

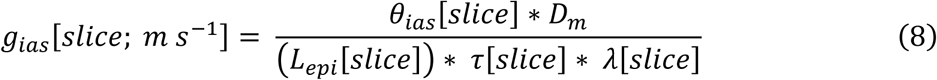

in which [slice] denotes per-slice average values. Equation 7 was then used to convert g_ias_ [slice; m s^−1^] into mol m^-2^ s^-1^ units.

### Display

Z-projections of tissue diameter (e.g. Fig. 2G-I) were calculated with FIJI/BoneJ’s Thickness tool, then Z-projected with FIJI’s “Average Z-project” tool and assigned an LUT (Doube et al., 2010). 3D images for display (e.g. Fig. 2K-V) were created with ImageJ/3DScript (Schmid et al., 2019), and stacks with diameter-labeled color masks (or monotone color mask in the case of airspace). Confocal images for Fig. 1A-B were produced using sorghum leaf tissue harvested fresh and stained with calcofluor white (CFW) diluted in PBS for approximately one hour under vacuum (Bidhendi et al., 2020), to provide alternative imagery of stomate and air crypt.

**Fig. 2:**
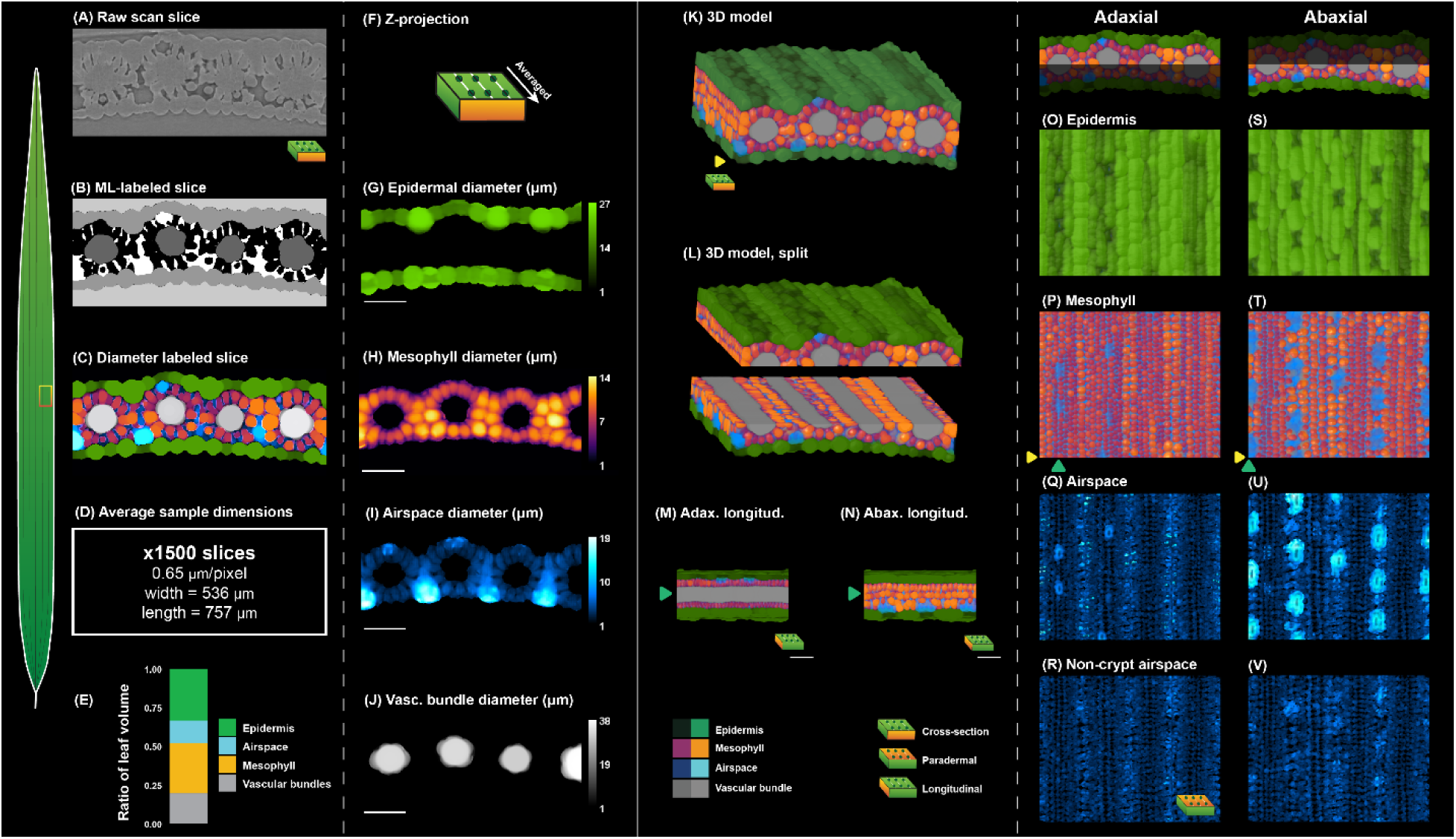
(A) Raw reconstructed single-slice microCT scan (B) machine-learning labeled single slice example (C) diameter-labeled epidermis, mesophyll, airspace, and vasculature ML classes (D) Dimensions and resolution for average leaf sample (E) volume ratios for each ML class across all samples (F) Z-projection process used for G-J (G-J) average z-projections of diameters for each ML class, from 0.8 mm long leaf section (K) 3D representation of Sorghum leaf (L) Panel K split between adaxial and abaxial regions (M) longitudinal view of adaxial stomates and sub-stomatal crypts (N) longitudinal view of abaxial stomates and sub-stomatal crypts (O-R) Paradermal adaxial epidermis, airspace/mesophyll, and diameter-labeled airspace; airspace divided into full airspace and airspace with stomate-associated air (crypts) filtered out (S-V) Paradermal abaxial epidermis, mesophyll, and diameter-labeled airspace; airspace divided into full airspace and airspace with stomate-associated air (crypts) filtered out.

### Statistical analyses

In all cases, data from multiple sub-samples within a leaf were averaged to generate means of a trait for a given leaf. To compare traits (e.g. stomatal density, diameters, conductance traits, etc.) between groups (AD v AB, or WT v EPF_syn_), a mixed-effects model was used from R package nlme (Pinheiro et al., 2025), to test for the fixed effects of leaf surface (adaxial, abaxial), genotype (WT, EPF_syn_), and the interaction of surface and genotype. Least-squares means post-hoc analyses were carried out with the lsmeans function from the lsmeans package in R (Lenth, 2016). To test genotypic differences within Z-depth curves, depth data was divided into different regions based on distance through the airspace, and then the means of each region were compared between genotypes via student’s t-test in R. For traits where a mixed-effects model was not appropriate to test for a genotype effect (e.g. absent file frequency, which occurred only on the adaxial surface), a student’s t-test was performed.

## RESULTS

### Full sample tissue ratios, mesophyll surface area and airspace connectivity

Airspace, mesophyll, vasculature and epidermises accounted for ∼15%, ∼33%, ∼19%, and ∼33% of total leaf volume, respectively (Fig. 2E). Mean values for these traits did not differ significantly between genotypes. Leaf porosity (θ) calculated from airspace and mesophyll volumes (Formula 1) was 31%.

Connectivity (κ_V_) was measured as the ratio between the volume of the largest airspace particle in the sample and the sum of all airspace particle volumes in the sample. On average, 99.3% of the leaf airspace was connected to the atmosphere, meaning that very few airspaces detected within the leaves were disconnected. Major veins (e.g. veins with sclerenchyma above *and* below the vascular bundle) divided the leaf airspace into two distinct and disconnected airspace regions lateral to the major vein (Supp. fig. 1), thus stacks were cropped to remove the regions associated with sclarified major veins prior to κ_V_ calculations. Small regions of disconnected airspace were typically scattered throughout the airspace, with no obvious localization (Supp. fig. 2). Unbroken pathways connected same-side sub-stomatal crypts to one another (e.g. movement between adjacent adaxial or abaxial stomatal files) and between the two sides (e.g. adaxial to abaxial). However, these pathways included thin air channels (<2 µm diameter) and tortuous regions (Supp. fig. 4), indicating low CO_2_ diffusion beyond sub-stomatal crypts.

### Stomatal density, dimensions, and patterning

Stomatal density was significantly higher on the abaxial surface in both WT (+85%) and EPF_syn_ (+150%) (Fig. 3A). Abaxial stomatal complexes were significantly longer (WT +20%; EPF_syn_ +16%), wider (WT +16%; EPF_syn_ +11%), and larger in area (WT +40%; EPF_syn_ +30%) compared to adaxial stomatal complexes in both genotypes (Fig. 3D, F). Stomatal frequency along a file of cells was significantly higher (WT +79%; EPF_syn_ +162%) on the abaxial surface compared to adaxial surface (Fig. 3B).

**Fig. 3:**
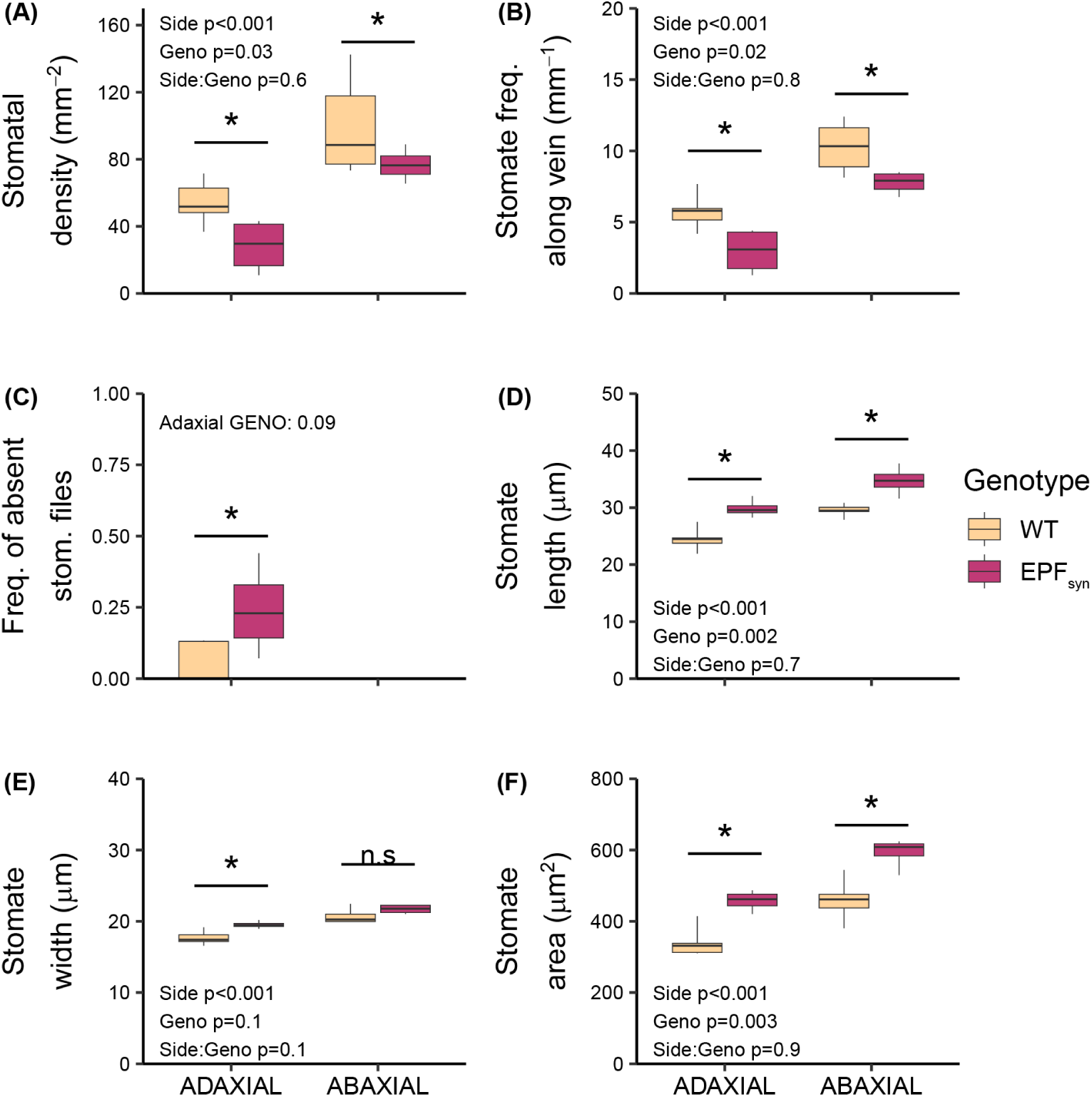
(A) Stomatal density (SD) is significantly higher on the abaxial side compared to adaxial. SD is significantly lower on both adaxial and abaxial sides in EPF_syn_. (B) Stomatal frequency along lengths of veins (i.e. number of stomates within a single stomatal file along a given length of vein) is higher on the abaxial side compared to the adaxial side in both genotypes, and is significantly reduced on both sides in EPF_syn_ compared to WT (C) the rate of “absent” stomatal files (i.e. vascular bundle lacking associated adaxial or abaxial stomatal file) was always zero on the abaxial side; there was significantly more absent stomatal files in EPF_syn_ compared to WT, but only on the adaxial side. (D) Length of the stomatal complex is higher on the abaxial side compared to adaxial in both genotypes, and is higher on both sides in EPF_syn_ (E) Stomate width is significantly higher on the abaxial side compared to adaxial in both genotypes, and is increased on the adaxial side in EPF_syn_ (F) Area of the stomatal complex is higher on the abaxial side compared to adaxial in both genotypes, and is higher on both sides in EPF_syn_ Asterisks indicate p<0.05. Error bars represent standard error.

The location of stomatal files in relation to underlying vascular bundles was different between adaxial and abaxial leaf surfaces (Fig. 4A-D). A significantly greater proportion of stomates were located over the vasculature on the adaxial surface (87%) than on the abaxial surface (3%). Adaxial stomates were generally located over the center of the vascular bundle or immediately adjacent to it. In contrast, abaxial stomates were generally located directly between pairs of vascular bundles (Fig. 4D-E).

**Fig. 4:**
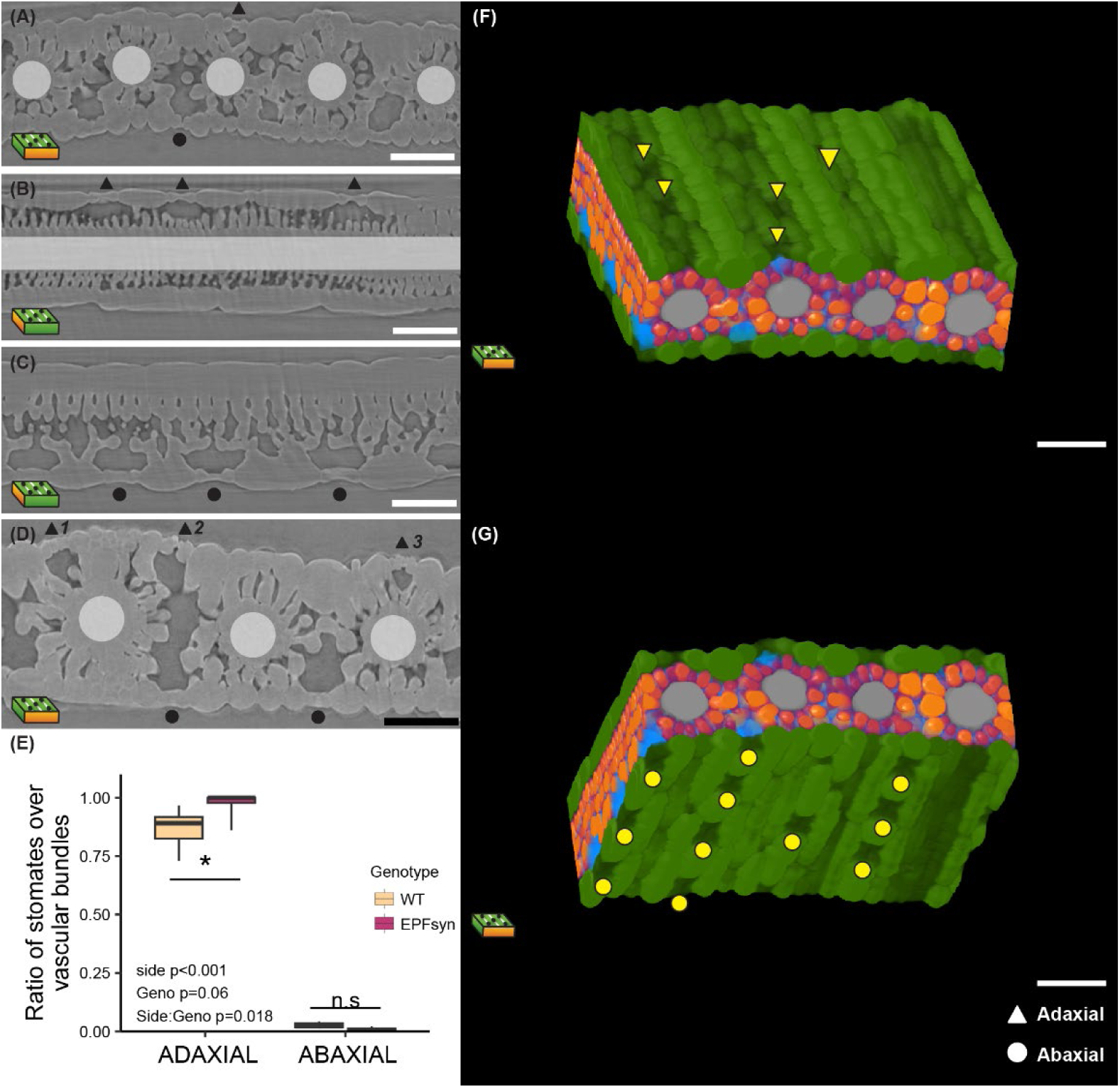
Different views of a sorghum leaf, showing relationships between adaxial and abaxial stomates to vascular bundles. Vascular bundles indicated by opaque white boxes and bar. Adaxial stomates indicated by triangle symbol and abaxial stomates indicated by circle symbol. (A) Cross-section view of sorghum leaf and stomates (B) Longitudinal view of sorghum leaf and stomates, sliced through adaxial stomatal file (C) Longitudinal view of sorghum leaf and stomates, sliced through abaxial stomatal file (D) 1: Adaxial stomate overlying vascular bundle but towards the periphery of the bundle. 2: Adaxial stomate NOT overlying the vascular bundle, uncommon. 3: Adaxial stomate overlying vascular bundle in the center of the bundle. (E) The ratio between the number of stomates overlying the vascular bundle to total number of stomates. Significantly higher on the adaxial side compared to the abaxial side, and significantly higher on the adaxial side in EPF_syn_ compared to WT. Asterisks indicate p<0.05. Error bars represent standard error. (F) 3D cartoon of adaxial leaf surface and internal anatomy, triangles indicate adaxial stomates (G) 3D cartoon of abaxial leaf surface and internal anatomy, triangles indicate adaxial stomates.

### Airspace traits and CO_2_ conductance

The abaxial airspace region had both greater overall bulk airspace diameter (crypt and non-crypt airspace; WT +84%; EPF_syn_ +109%; Fig. 6A) and greater non-crypt airspace diameter then the adaxial airspace region (WT +24%; EPF_syn_ +28%; Fig. 6B). Similarly, the abaxial region of the leaf had greater θ (WT +37%; EPF_syn_ +44%; Fig. 6G). Abaxial crypts were larger in all dimensions relative to adaxial crypts, in both genotypes. Abaxial crypts were more voluminous (WT +622%; EPF_syn_ +818%), longer (WT +98%; EPF_syn_ +98%), wider (WT +114%; EPF_syn_ +135%), and deeper (WT +227%; EPF_syn_ +290%) than adaxial crypts (Fig. 5D-G).

**Fig. 5:**
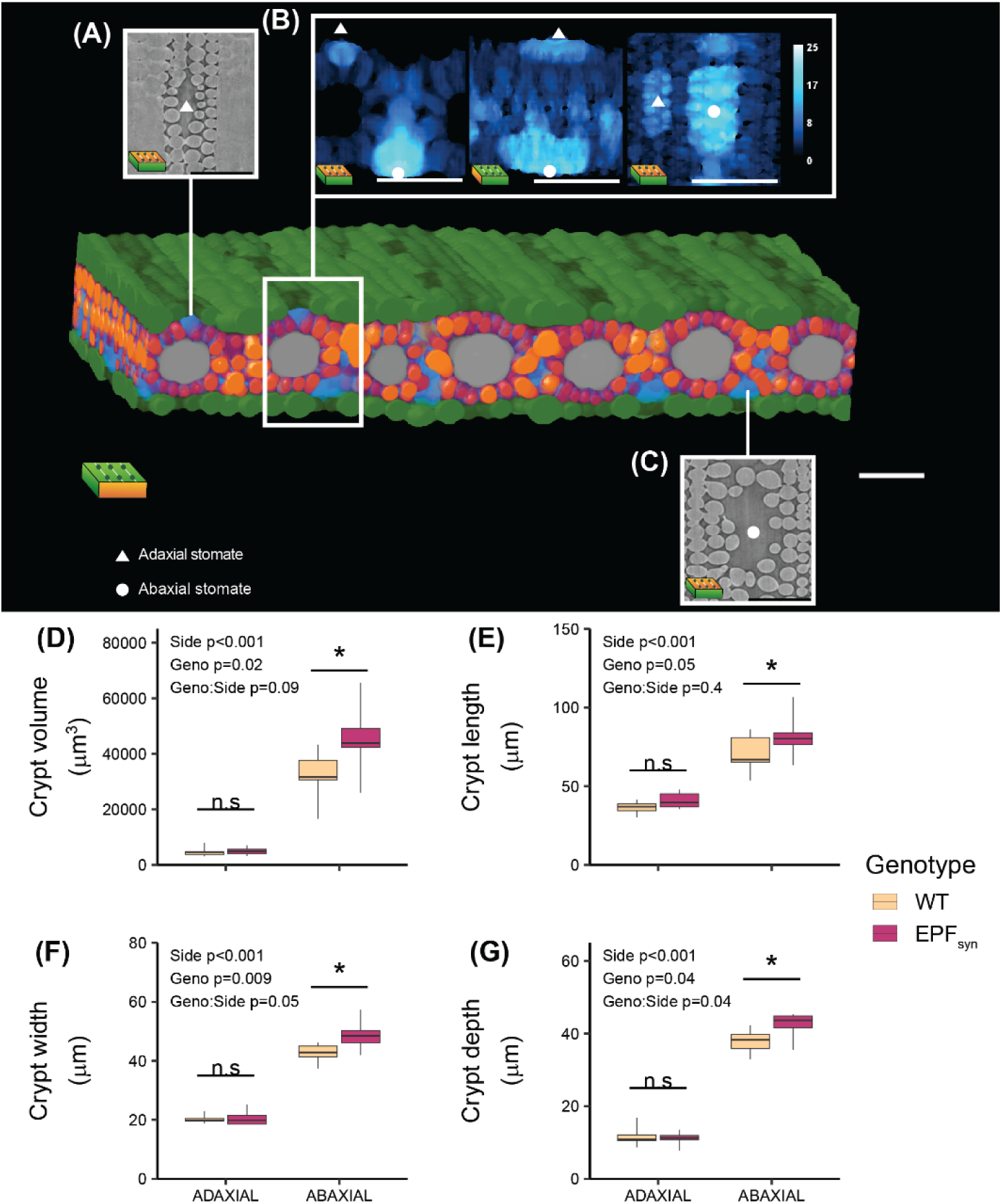
(A) Raw paradermal microCT scan of adaxial sub-stomatal air crypt (B) average z-projection of airspace diameter labeled adaxial sub-stomatal air crypt and abaxial sub-stomatal air crypt from different viewing angles (C) raw paradermal microCT scan of abaxial sub-stomatal air crypt (D-G) adaxial and abaxial sub-stomatal crypt dimensions in wildtype and EPF_syn_. Crypt volume, length, width and depth were all significantly higher on the abaxial side. Abaxial crypts were significantly larger in all dimensions in EPF_syn_. Asterisks indicate p<0.05. Error bars represent standard error.

**Fig. 6:**
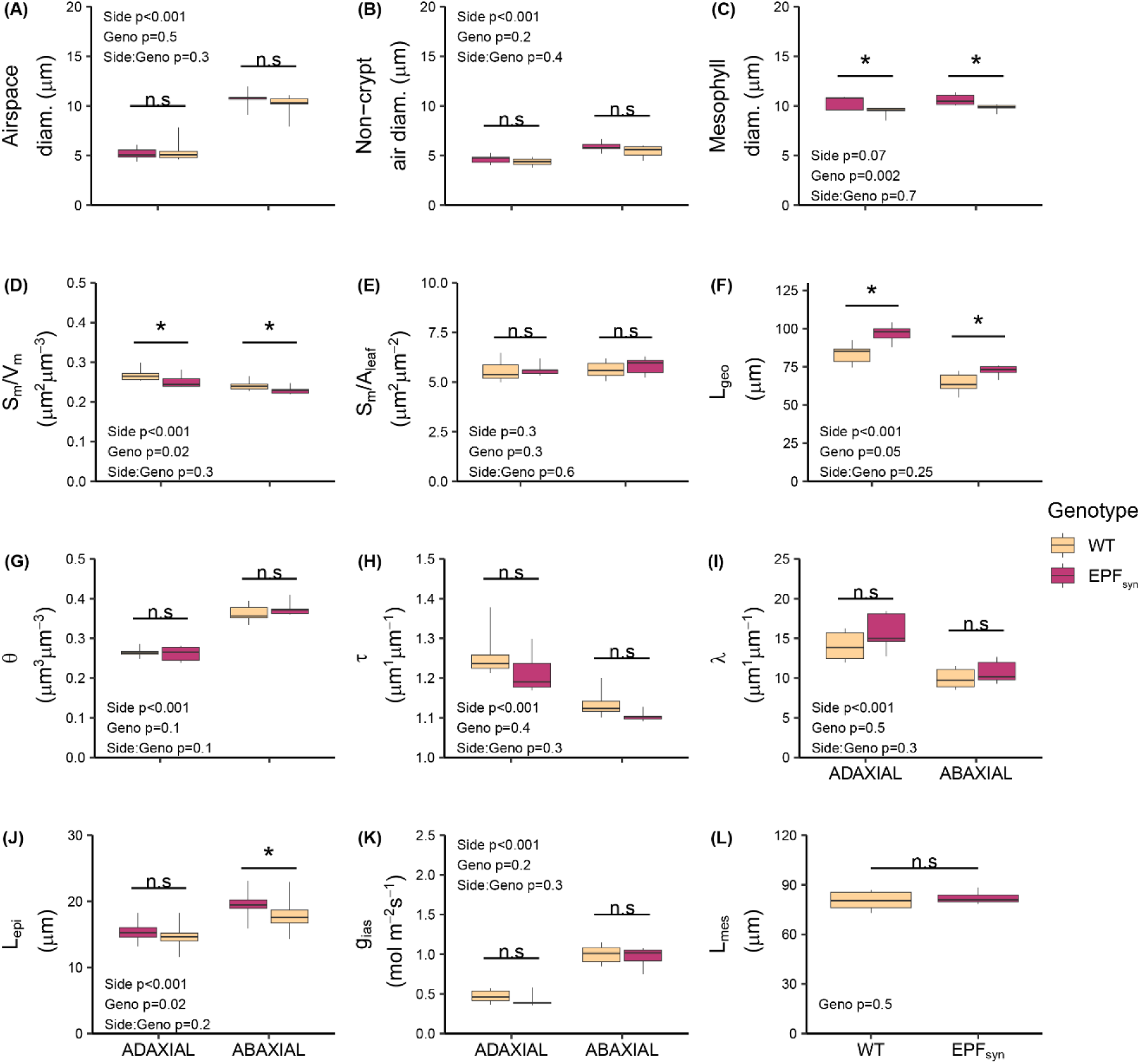
Airspace, mesophyll, and g_ias_ traits measured between adaxial and abaxial regions of the leaf in the two genotypes. (A) Adaxial and abaxial average airspace diameter (crypt and non-crypt airspaces analyzed together) (B) average non-crypt airspace diameter in adaxial and abaxial regions (C) overall mesophyll diameter (D) average ratio between the area of airspace in contact with mesophyll (S_m_) and the sum of airspace and mesophyll volumes (V_m_) (E) average ratio between S_m_ and sample epidermal area (A_leaf_) (F) average distance between stomate and any point along the surface of the mesophyll in the adaxial or abaxial regions (L_geo_) (G) average porosity (θ) calculated with formula 1 (H) average airspace tortuosity (τ) calculated with formula 4 (I) average airspace lateral path lengthening (λ) calculated with formula 5 (J) average distance from adaxial or abaxial epidermis to any point along the mesophyll (L_epi_) (K) average airspace conductance to CO_2_ (g_ias_) calculated with formulas 6 and 7 (L) average leaf thickness, without epidermises (L_mes_). F-L were measured at the region where airspace and mesophyll meet. Asterisks indicate p<0.05. Error bars represent standard error between biological replicates.

The abaxial side had greater S_m,V_ (WT +11%; EPF_syn_ +10%, Fig. 6D), but there was no significant difference in S_m,LA_ between sides (Fig. 6E). The abaxial side also had lower L_geo_ (WT -23%; EPF_syn_ -25%; Fig. 6F) and greater L_epi_ (WT +21%; EPF_syn_ +26%; Fig. 6J). With lower τ (WT -10%; EPF_syn_ -9%; Fig. 6H) and lower λ (WT -29%; EPF_syn_ -32%; Fig. 6I), abaxial g_ias_ was significantly greater than adaxial g_ias_ in both genotypes (WT +42%; EPF_syn_ +39%; Fig. 6K).

### EPF_syn_

SD was significantly lower in EPF_syn_ relative to WT on both the adaxial (-42%) and abaxial (-22%) surfaces (Fig. 3A). EPF_syn_ expression led to a significant reduction in adaxial (-49%) and abaxial (-24%) stomatal frequency within a cell file relative to WT (Fig. 3B). Stomatal complexes were significantly longer in EPF_syn_ relative to WT (Fig. 3D) on both adaxial (+22%) and abaxial surfaces (+18%). Similarly, stomatal complex area was significantly greater in EPF_syn_ (Fig. 3F) on both adaxial (+34%) and abaxial surfaces (+24%).

Compared to WT, EPF_syn_ plants had greater mesophyll cell diameter (AD +11%; AB +10%; Fig. 6C) and decreased S_m,V_ (AD -6%; AB -6%; Figs. 6D). Across the vertical profile, the mesophyll cell diameter of EPF_syn_ compared to WT was significantly greater in both the deeper regions of the abaxial side (+7%) and regions of the abaxial side close to the epidermis (+9%). Overall airspace diameter and non-crypt airspace diameter were not significantly different when analyzed in bulk (Fig. 6A, 6B). Diameter of *non-crypt* airspaces (e.g. inter-connecting air pathways) for EPF_syn_ compared to WT was significantly greater in the lower layers of the abaxial region (+18%) and marginally greater in the upper regions of the adaxial side (+11%; Fig. 7B).

**Fig. 7:**
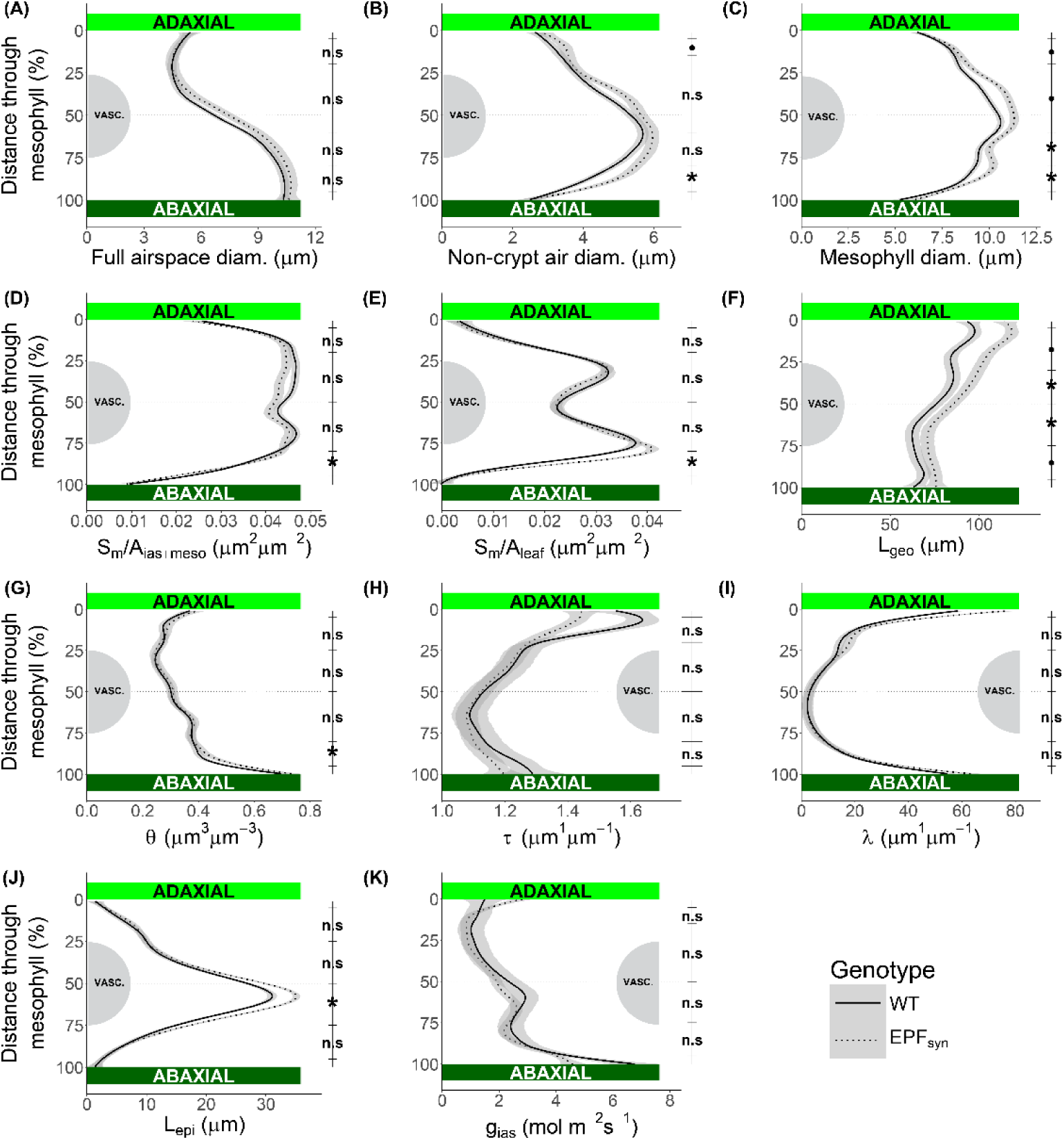
Airspace, mesophyll, and g_ias_ traits measured on a slice-by-slice basis, moving from the adaxial epidermis to the abaxial epidermis (1-100%). Gray half-circles represent approximate location of vascular bundles. Average slice-by-slice values shown for: (A) Full airspace diameter (B) Non-crypt airspace diameter (C) Mesophyll diameter (D) average ratio between the area of airspace in contact with mesophyll (S_m_) and the sum of airspace and mesophyll volumes (V_m_) (E) average ratio between S_m_ and sample epidermal area (A_leaf_) (F) distance from stomate to any point along the mesophyll (L_geo_) (G) porosity (θ) calculated with formula 1 (H) tortuosity (τ) calculated with formula 4 (I) lateral path lengthening (λ) calculated with formula 5 (J) distance from epidermis (L_epi_) (K) airspace conductance to CO_2_ (g_ias_) calculated with formula 8. Translucent gray envelopes represent SE between biological replicates of a genotype. Symbols indicate significant difference between WT and EPF_syn_ in the indicated region of leaf depth: .<0.1 *<0.05.

Despite no shift in overall airspace diameter, there was a general genotype by leaf surface interaction effect on crypt size, such that pairwise tests showed significantly greater dimensions. EPF_syn_ crypts were 43% more voluminous, 20% longer, 16% wider, and 11% deeper than WT on the abaxial side, but not the adaxial side (Fig. 5D-G). This indicates that while EPF_syn_ expression reduced the density of sub-stomatal crypts (via SD reduction), this did not reduce overall airspace diameter, due to the increase in crypt sizes.

EPF_syn_ θ was unaffected when analyzed in bulk (Fig. 6G), however EPF_syn_ θ was marginally greater than WT in a region of the abaxial side close to the epidermis (+8%; Fig. 7G) when analyzed in detail across the leaf vertical profile. Across the airspace geometric traits when analyzed in bulk or in detailed vertical profile, only L_geo_ and L_epi_ were affected in EPF_syn_. In bulk, EPF_syn_ had greater L_geo_ on both sides (AD +15%; AB +12%; Figs. 6F, 7F) and greater abaxial L_epi_ (AB +10%; Figs. 6J, 7J). τ and λ were unaffected in EPF_syn_ in both bulk and vertical profile analyses (Figs. 6H-I, 7H-I). As such, g_ias_ was unaffected overall in EPF_syn_ when analyzed in bulk and in vertical profile (Figs. 6K, 7K).

## DISCUSSION

Here, it was successfully demonstrated that microCT methods could be used to assess coordination in the patterning of stomata, airspaces, mesophyll and vascular bundles in the model C4 grass sorghum, expanding the list of tractable model taxa for future investigations of the relationships between leaf structure and function. Additionally, the resulting data did not support any of the hypotheses developed based on previously available information. First, stomates and associated IAS volumes were not consistently localized solely between vascular bundles as has been previously reported in other species and observed in published cross-section microscopy images of non-sorghum species (Long et al., 1989; Lundgren et al., 2019; McKown and Bergmann, 2020; Schuler et al., 2018; Sonawane et al., 2021; Stebbins and Shah, 1960). Instead, there were numerous instances of stomata positioned over rather than between vascular bundles on the adaxial surface. Those adaxial stomates represent up to 95% of total adaxial stomata (Fig. 4). In contrast, abaxial stomates and associated IAS were nearly always localized between veins. Second, the development of stomates and associated air crypts over vascular bundles resulted in far smaller adaxial air crypts compared to abaxial, which led to the g_ias_ of the adaxial surface being disproportionately lower than would be predicted from the ratio of SD between the two leaf surfaces. However, despite the much smaller crypts the adaxial IAS contributed significantly to overall g_ias_ and could play important physiological roles separate from abaxial stomates, which should be investigated in future work. Third, θ and g_ias_ were not significantly different between low-SD and wildtype sorghum, because there was plasticity in other airspace traits that counteracted one another in a leaf surface-specific manner. In addition, the results reveal plasticity in the internal airspaces of C4 crops engineered to have low SD, which circumvent expected changes in g_ias_ that would otherwise impede efforts to improve iWUE.

### Advances in analyzing leaf internal airspace

High-resolution microCT produces large gray-scale stacks of images, requiring tissue labeling to enable downstream analyses (Théroux-Rancourt et al., 2020). Machine learning (ML) powered segmentation avoids the necessity of hand labeling different tissues across the hundreds or thousands of slices within a sample, a time-consuming process which previously limited experiments to a small number of slices and/or samples (Théroux-Rancourt et al., 2020). Furthermore, due to the low availability of machines capable of high-resolution microCT scanning (e.g. large synchrotron radiation facilities), the utilization of this tool has only begun to accelerate in the plant sciences (Piovesan et al., 2021). As such, certain functional groups of species have yet to be analyzed in high-resolution 3D, such as C4 species (Théroux-Rancourt et al., 2021).

Here, a sorghum-specific ML pipeline was trained to streamline the segmentation process, using 20 training slices distributed over the whole sample set, instead of building a training dataset for each sample or labeling all slices by hand (Théroux-Rancourt et al., 2020). The analysis was furthered by investigating the vertical profiles of interior airspace traits (e.g. diffusive path length) and g_ias_ across the leaf depth, allowing for more detailed analyses (e.g. slice-by-slice vertical profile, adaxial versus abaxial) than previous bulk-leaf measurements (e.g. Harwood et al., 2021). Additionally, to accurately study sub-stomatal crypts, a method was devised to extract and analyze these air spaces, offering one of the first quantifications of crypts in three dimensions. As such, this study provides a new in-depth study of sub-stomatal air crypts, and one of the first high-resolution investigations into the C4 leaf interior.

### Asymmetry in stomatal patterning and sub-stomatal crypt morphology between adaxial and abaxial sides

High-resolution microCT revealed that 99.3% of leaf airspace was connected, with few isolated airspace regions (Supp. figs. 1, 2). Within this airspace, voluminous sub-stomatal air crypts were interconnected via a complex network of thin (approx. 5 µm average diameter; Fig. 6B) and tortuous air channels formed by the complex geometry of the mesophyll, bundle sheath, and veins (Supp. fig. 3; Supp. video 1). Stomates co-develop with underlying mesophyll: expansion and deformation of the underlying mesophyll cells create the sub-stomatal air crypt (Baillie and Fleming, 2020). However, there has been little detailed research on the sub-stomatal air crypts themselves, potentially as a consequence of the limitations of two-dimensional microscopy (Théroux-Rancourt et al., 2023).

There was a striking difference in the internal anatomy and in size between adaxial and abaxial crypts (Figs. 4, 5). Adaxial sub-stomatal crypts were surrounded primarily by smaller mesophyll cells located at the apex of the radial mesophyll bundle, while abaxial crypts were surrounded by larger radial mesophyll cells located between vascular bundles (Fig. 2). Adaxial crypts were smaller in all dimensions (volume, length, width, depth), with the extent of the adaxial crypt truncated by underlying vascular bundles (Figs. 4, 5). The contrasting location of abaxial stomatal files between vascular bundles allowed abaxial crypts to extend in space much more than their adaxial counterparts (Figs. 4, 5). It is possible that small adaxial crypts are a direct result of the physical location of adaxial stomata over vascular bundles, with less space to expand between the epidermis and veins.

Contrary to expectation, in wildtype plants, 87% of adaxial stomatal files were located over vascular bundles (Fig. 4). The position of the stomates were often centered directly over the vascular bundle (example adaxial stomata #3, Fig. 4D), sometimes over the edge of the vascular bundle (example adaxial stomata #1, Fig. 4D), but rarely in a central location between vascular bundles (example adaxial stomata #2, Fig. 4D) to mirror the consistent position of abaxial stomatal files (Fig. 4D). This observation runs counter to the widely held assumption that stomates develop without overlapping the vasculature (Baresch et al., 2018; Esau, 1965; McKown and Bergmann, 2020) and identifies a distinction between the two surfaces of the leaf that could be functionally significant. Micrographs of sorghum leaves in an independent study corroborate that adaxial stomata are found over vascular bundles, even if that study did not discuss the arrangement (de Oliveira et al., 2022).

The position of adaxial stomata over vascular bundles may be a consequence of competition with other cell types for the region between vasculature on the adaxial epidermis. In this region, sorghum leaves develop rows of bulliform cells, which are an important drought tolerance-related cell type that enable leaf curling under drought stress (Seleiman et al., 2021). Development of adaxial bulliform cells between the vascular bundles may “push” adaxial stomatal files to the unoccupied regions directly overlying vascular bundles. However, despite the surface competition for epidermal space between stomata and bulliform cells, and the internal competition between crypt airspace and mesophyll, stomata are still present on the adaxial surface. This may indicate that adaxial stomata and underlying air crypts are still vital in providing CO_2_ to the leaf interior, despite small crypts and lower SD on the adaxial side. Furthermore, analysis on the paths connecting airspaces between the two sides indicates that although there are unbroken paths through the airspace connecting adaxial and abaxial airspaces, these paths are tortuous and pass through very thin airspaces (<2 µm diameter; Supp. fig. 4). Thus, adaxial stomata most likely provide CO_2_ to a distinct set of mesophyll cells separate from mesophyll cells on the abaxial side of the leaf, with low gaseous flow between the two sides (Long et al., 1989).

### CO_2_ conductance asymmetry in the leaf airspace

Here, the Sorghum leaf airspace had high asymmetry in traits between leaf sides which factor into the airspace’s conductance to CO_2_ (g_ias_). The different morphology of the adaxial region led to lower g_ias_ compared to the abaxial region: lower SD, lower θ, higher τ, and higher λ (Figs. 6-9). Higher λ occurred in the adaxial airspace compared to the abaxial side as a result of lower SD on the adaxial side and the close distances of mesophyll to the epidermis in the adaxial region of the leaf (lower average L_epi_; Fig. 6J). τ was highest in the adaxial airspace, and in the diagonally seated regions of airspace around the vasculature, generally laterally situated relative to adaxial stomatal files (Figs. 6H, 7H, 8I), indicating thin and complex airpaths directly outside of sub-stomatal crypts. The most likely driver behind g_ias_-trait asymmetry between adaxial and abaxial sides is the much smaller sub-stomatal crypts associated with adaxial stomata. This asymmetry in crypt sizes drives increased τ and λ in the adaxial region. Denser and more difficult to navigate mesophyll associated with smaller crypts increases τ, while adaxial mesophyll associated with adaxial crypts are closer to the surface of the leaf than their abaxial counterparts which increases λ. Exploring the functional consequences of such an arrangement should be the subject of future studies.

**Fig. 8:**
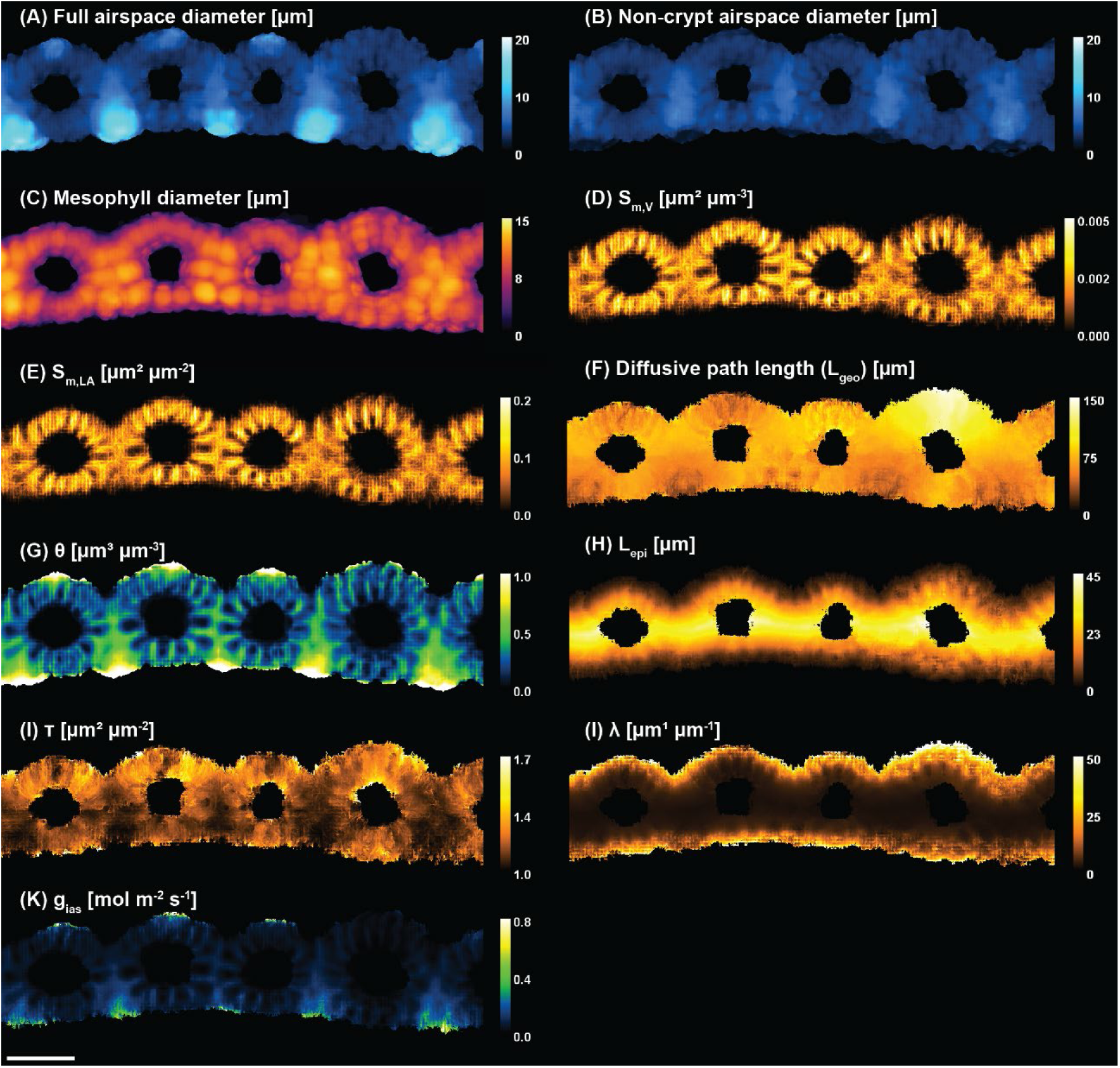
Average z-projections of traits plotted in figures 5 and 6. The same wildtype sub-sample is shown across panels. (A) Airspace diameter (crypt and non-crypt airspaces) (B) non-crypt airspace diameter (C) average mesophyll diameter (D) ratio between the area of airspace in contact with mesophyll (S_m_) and the sum of airspace and mesophyll volumes (V_m_) (E) ratio between S_m_ and sample area (A_leaf_) (F) distance between stomate and any point along the surface of the mesophyll (L_geo_) (G) average porosity (θ) (H) airspace tortuosity (τ) (I) lateral path lengthening (λ) (J) distance from epidermises to any point along the mesophyll (L_epi_) (K) average airspace conductance to CO_2_ (g_ias_).

**Fig. 9:**
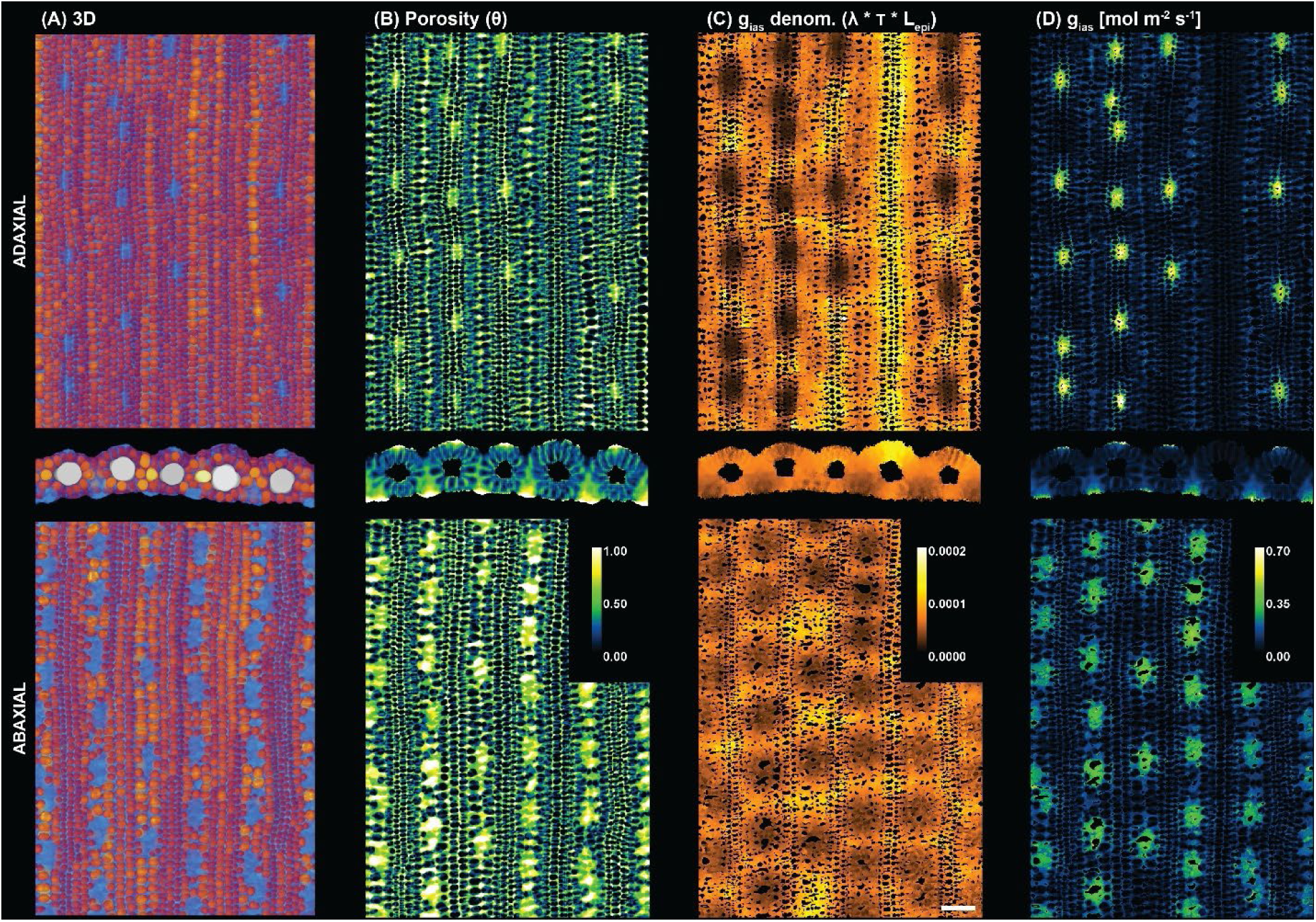
(A) 3D cartoons of adaxial, cross-section, and abaxial mesophyll and airspace (B) average z-projected porosity of adaxial, cross-section, and abaxial airspace regions of the leaf (C) average z-project of the product of lateral path lengthening (λ), tortuosity, (τ) and distance from epidermis (L_epi_) at the mesophyll/airspace contact boundary for adaxial, cross-sectional, and abaxial regions of the leaf (D) Average g_ias_ for adaxial, cross-sectional, and abaxial regions of the airspace, at the mesophyll/airspace contact boundary.

Inevitably, regions distant from stomates—between stomatal files or in a region of the leaf with patchy stomata, for example—had higher τ, high diffusive path length (L_geo_), or lower θ. Regions of very low g_ias_ were often located in the region located between vascular bundle and abaxial epidermis, or in the regions between abaxial stomatal files, and over vascular bundles which lacked adaxial stomatal files (Figs. 8K, 9D). Low g_ias_ also occurred between stomates *within* a file. Therefore, a potential consequence of the well-organized stomatal file system are predictable “dead zone” regions of airspace with low potential gas exchange contributions (Figs. 8K, 9D) (Earles et al., 2018; Théroux-Rancourt et al., 2023). Colocalization of high τ and high λ contributed to g_ias_-trait heterogeneity and could have implications for strong [CO_2_] gradients within the leaf. Similarly, Earles et al. found colocalization of g_ias_-traits and substantial [CO_2_] gradients in C3 and CAM bromeliads (Earles et al., 2018). It is notable that the CO_2_ concentrating mechanism of C4 photosynthesis allows for greater draw-down of [CO_2_] in mesophyll airspace, without imposing limitations on photosynthetic carbon gain. In fact, this is the basis for greater iWUE in C4 than C3 species. So, it may be valuable to compare the tolerance for dead zones in C3 species too.

Overall, g_ias_ calculated here (0.38-0.62 mol m^-2^ s^-1^) falls below reported C3 g_ias_ values (Earles et al., 2018; Harwood et al., 2021; Théroux-Rancourt et al., 2023) but above CAM values (Earles et al., 2018). This may indicate a lower threshold for g_ias_ required for photosynthesis in C4 compared to C3, due to the CCM present in C4 species. Further research into the airspace CO_2_ conductances across C3, C4, and CAM species is required to further understand the relationships between photosynthetic types and g_ias_. Until recently, the data required to calculate airspace conductance was limited to a few optical microscopy slices or prone to imaging and embedding artifacts, limiting the accuracy of the estimated values (Earles et al., 2018; Théroux-Rancourt et al., 2020). Here, a proof of concept is provided for studying g_ias_ in a C4 species with microCT, which is a method that can address key knowledge gaps moving forward.

### Stomatal patterning and airspace conductance in a reduced stomatal density transgenic (EPF_syn_)

As was found previously by Ferguson et al. (2024), the expression of a synthetic EPF protein (EPF_syn_) drove a significant reduction in the stomatal densities of the Sorghum leaf epidermises. EPF_syn_ also had larger stomatal complexes on both adaxial and abaxial surfaces than wild-type (Fig. 3D, F). Interestingly, EPF_syn_ abaxial sub-stomatal crypts were larger than WT air crypts in length, width, and volume (Fig. 5). With a decrease in SD, it appears as if abaxial crypts expanded further in space. In a two-dimensional analysis, de Oliveira et al. (2023) found that stomata-associated airspaces in sorghum increased in area under drought stress, indicating plasticity in airspace anatomy in sorghum (de Oliveira et al., 2023). Here, plasticity in three-dimensional airspace anatomy was found associated with SD-reduction. Furthermore, adaxial crypts appeared to be less plastic than abaxial crypts in their response to reduced SD (Fig. 5), introducing further asymmetry between the adaxial and abaxial regions of the leaf in EPF_syn_. Future work addressing differences in adaxial/abaxial crypt plasticity will offer further insights into the potential physiological differences between the two sides (e.g. if the crypts of one side are most responsive under drought conditions).

With reduced stomatal densities, one would expect g_ias_ to be concomitantly reduced, as the number of sub-stomatal crypts—in which g_ias_ is the most substantial (Fig. 9D)—are reduced at the same frequency as SD (e.g. a 1:1 relationship between stomate and crypt). However, g_ias_ was unaffected in EPF_syn_ despite the reduction in stomatal density and crypt density. In prior work on reduced-SD plants, a decrease in θ was concomitant with SD-reduction (Lundgren et al., 2019). However, no reduction in θ was found in EPF_syn_ sorghum, which could be attributed to the increase in abaxial crypt volume described above, compensating for the reduction in crypt numbers. Furthermore, while larger airspaces and crypts are associated with lower τ, low SD is associated with higher λ; these two responses were observed in EPF_syn_ though not statistically significant. These opposing responses in EPF_syn_ even each other when referring to Formula 7 and lead to g_ias_ being unaffected compared to WT, given the similar θ. Thus, EPF_syn_ maintained CO_2_ conductance compared to WT. This suggests that either the leaf is actively adjusting its internal anatomy to be optimized for a specific g_ias_, or that the developmental changes resulting from reduced SD do not influence gas diffusion in the internal airspace in a significant way, requiring future research to fully understand the relationships between SD-reduction severity and g_ias_. Future work should address the limits at which SD-reduction begins to significantly affect g_ias_ in a physiologically detrimental manner, and how it may interact with changes in stomatal aperture that are observed to compensate for the impact of lowering SD on stomatal conductance (Lunn et al., 2024).

## Conclusions

Here, a surprisingly large difference between adaxial and abaxial sub-stomatal crypts, both in size and in relation to mesophyll and underlying vasculature was discovered in *Sorghum bicolor*. The differences between adaxial/abaxial stomatal placement and associated air crypt dimensions raises further questions on the physiological implications of these differences, and the developmental signals creating these differences. Furthermore, there was high heterogeneity in airspace conductance traits across the leaf profile, such as high diffusive path length and high tortuosity in the adaxial airspace. As a consequence, the abaxial region of the leaf had higher airspace conductance potential. Furthermore, reduced stomatal density genotype EPF_syn_ had significantly larger abaxial sub-stomatal air cavities and significantly altered airspace traits while maintaining g_ias_ similar to WT. Future work should investigate the gas exchange contributions of the two leaf sides, as well as how crypt dimensions and stomatal placement interact with stomatal patterning and opening/closing. Such work will now be feasible given the technical advances in X-ray microCT reported here, along with new methods for assessing the development and structure-function relationships of stomata (Crawford et al., 2025; Hodge et al., 2025; Jain et al., 2021; Xie et al., 2021). In conclusion, a deeper understanding of adaxial/abaxial morphological differences, their physiological implications, and the inter-species variation in these traits can improve the ability to alter water use efficiency and gas exchange in key crop species.

## Acknowledgements

This research was supported by the DOE Office of Science, Office of Biological and Environmental Research (BER), grant numbers DE-SC0023160 and DE-SC0018277. This research used resources of the Advanced Photon Source, a US Department of Energy (DOE) Office of Science User Facility operated for the DOE Office of Science by Argonne National Laboratory under Contract No. DE-AC02-06CH11357. We would like to thank Dr. Devin Rippner from the US Department of Agriculture for helping with machine learning. We thank Pavel D. Shevchenko from Argonne National Laboratory for guidance on the Advanced Photon Source beamline. We thank Dr. Duncan Nall and Dr. Kingsley Boateng from the Core Facilities at the Institute of Genomic Biology for help with confocal analyses and software guidance. C.R.B. was supported by the Howard and Maryam Newman Plant Science Fund.

## Author contributions

A.D.B.L. and C.R.B. conceived and supervised the research, with contributions from J.M.F. and G.T.R. J.M.F. and C.R.B. collected the data. J.M.F., A.D.B.L., and G.T.R. developed methods and performed analyses of data. J.M.F., G.T.R., C.R.B., and A.D.B.L. wrote the manuscript.

## Data and code availability

The datasets used and coding implementation during the current study are publicly available from the online Illinois Databank at [DOI LINK TO ILLINOIS DATABANK, available upon acceptance] and GitHub repository [LINK TO GITHUB REPOSITORY, available upon acceptance].

---

**Supp. fig. 1:**
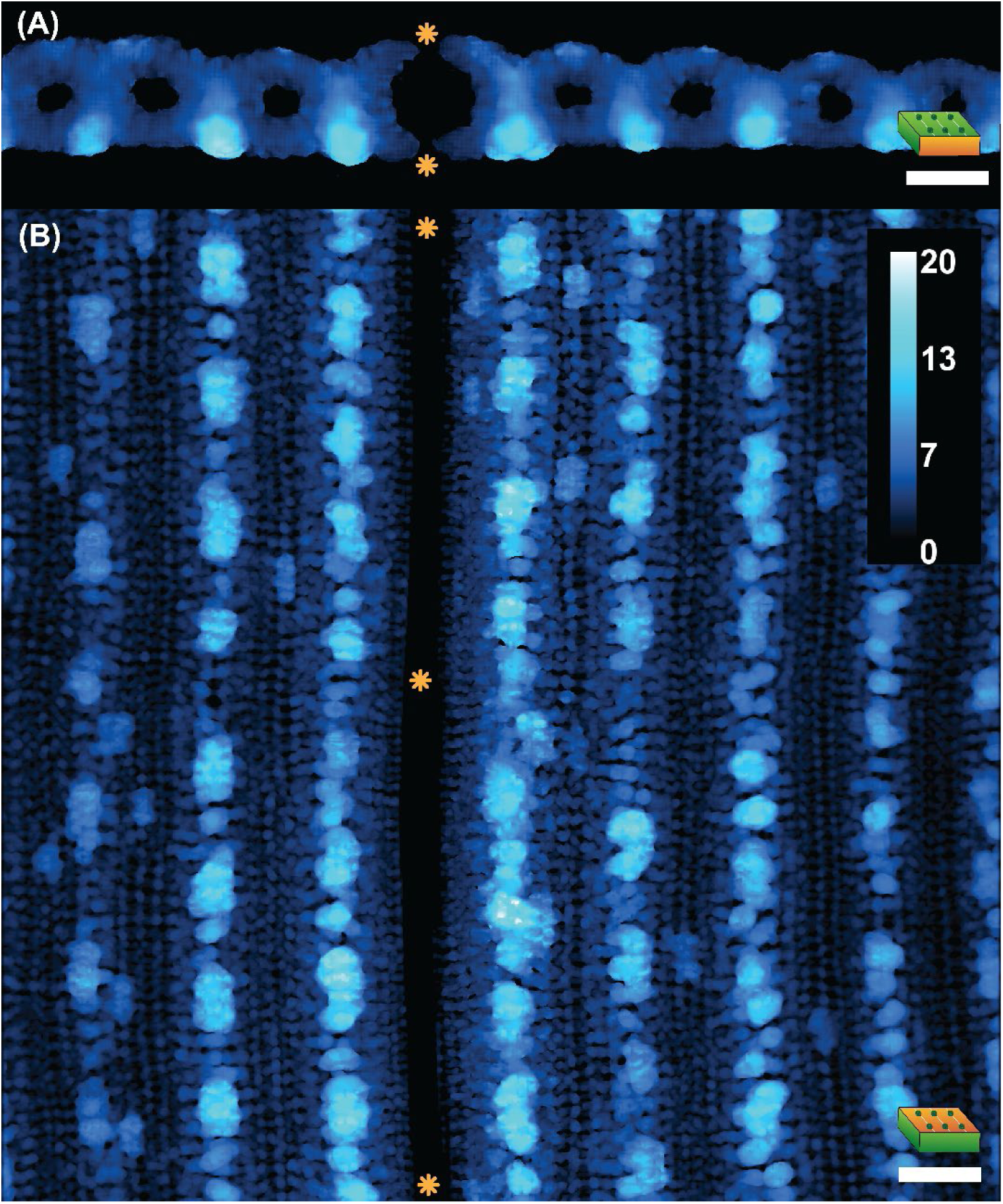
Average z-projections of airspace diameter (µm) for a sample with a heavily sclarified vascular bundle. Orange asterisks indicate heavily sclarified vasculature, which precludes formation of airspaces, dividing the airspace into two distinct regions lateral to the sclarified vascular bundle (A) cross-section z-projection (B) paradermal z-projection.

**Supp. fig. 2:**
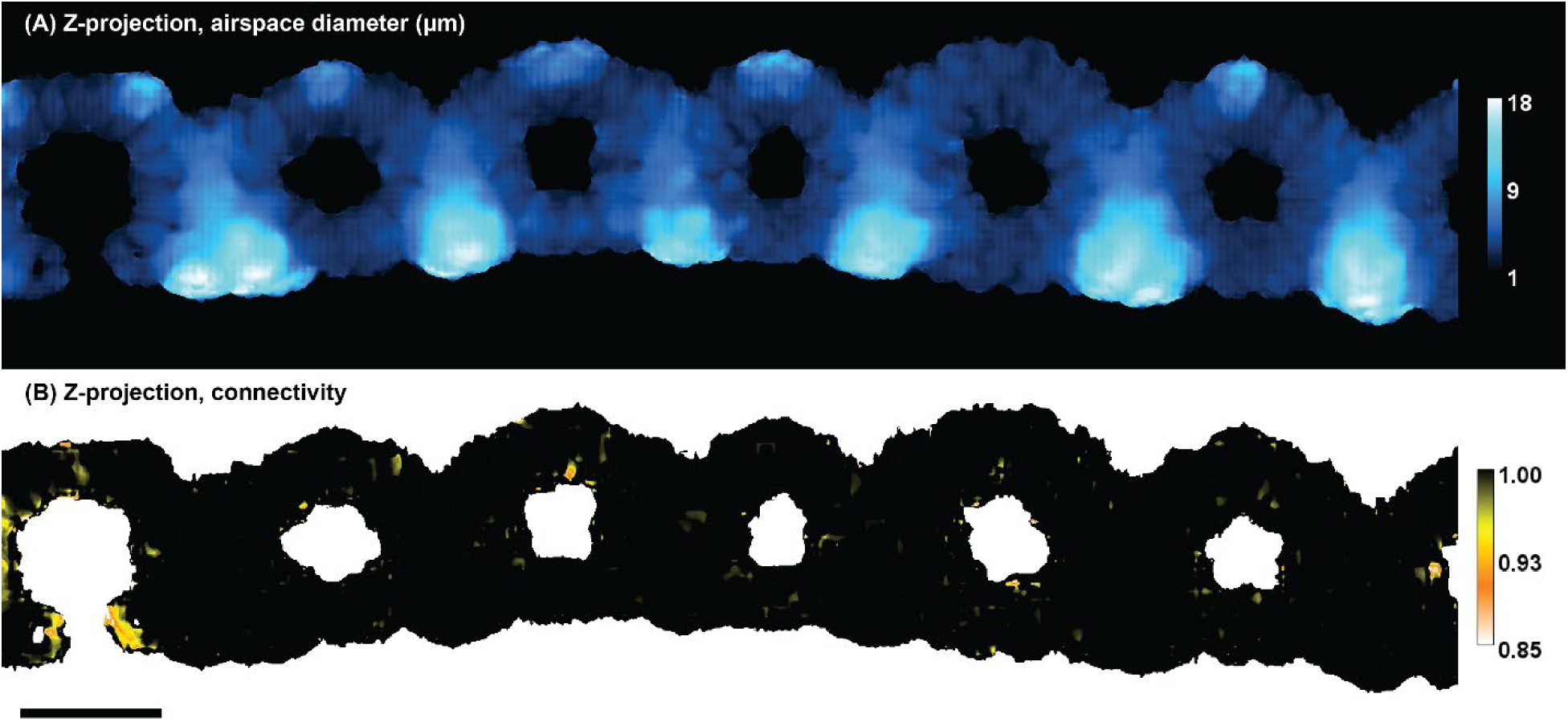
(A) Z-projected airspace diameter (B) Z-projected airspace connectivity, calculated from full airspace sum z-projection and z-projection of largest single airspace cavity in the sample. Regions with more regions of disconnected airspace are in warmer colors (e.g. <1.0).

**Supp. fig. 3:**
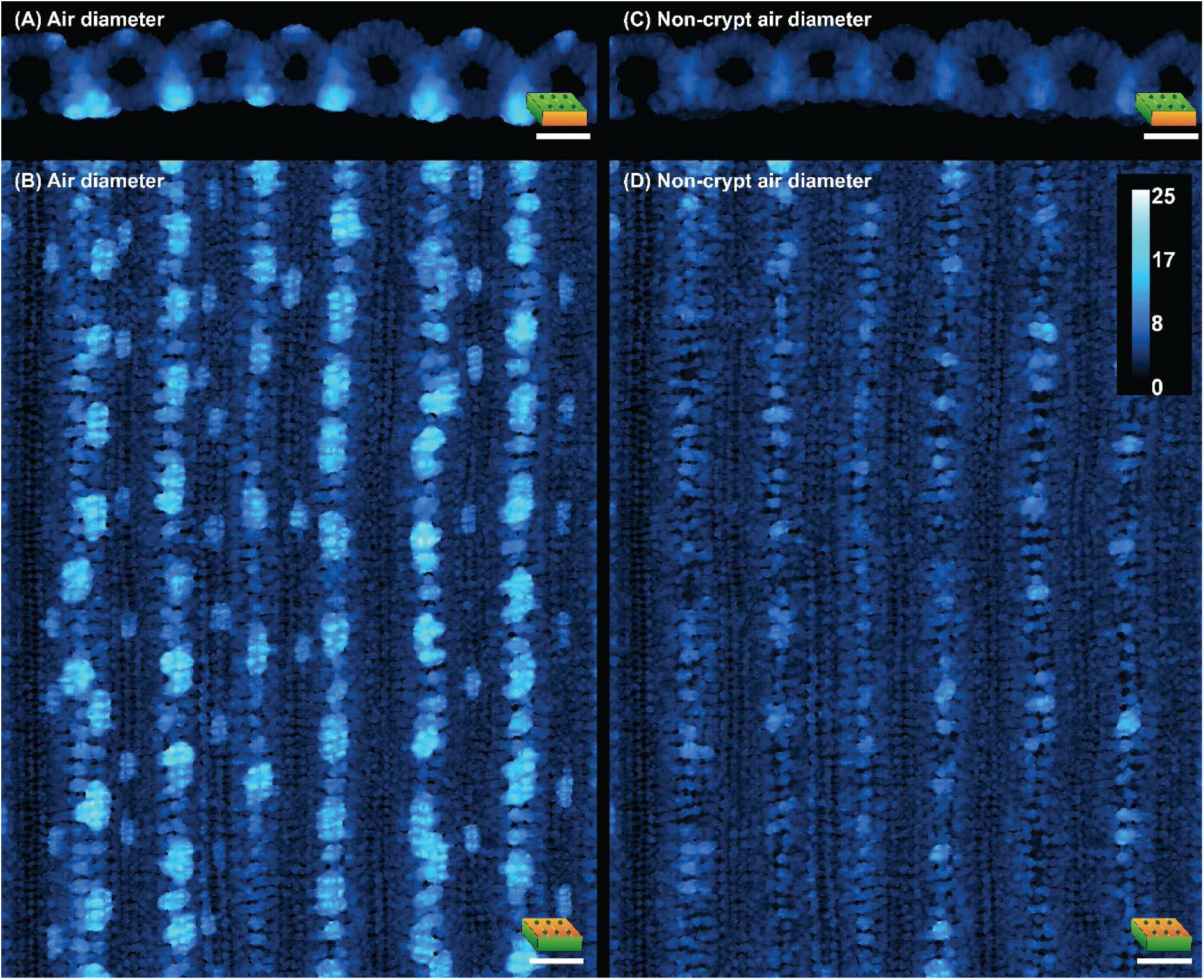
Average z-projections of airspace diameter for (A-B) entire airspace, including crypt and non-crypt airspace (C-D) Same airspace as A-B but with sub-stomatal air crypts removed.

**Supp. fig. 4:**
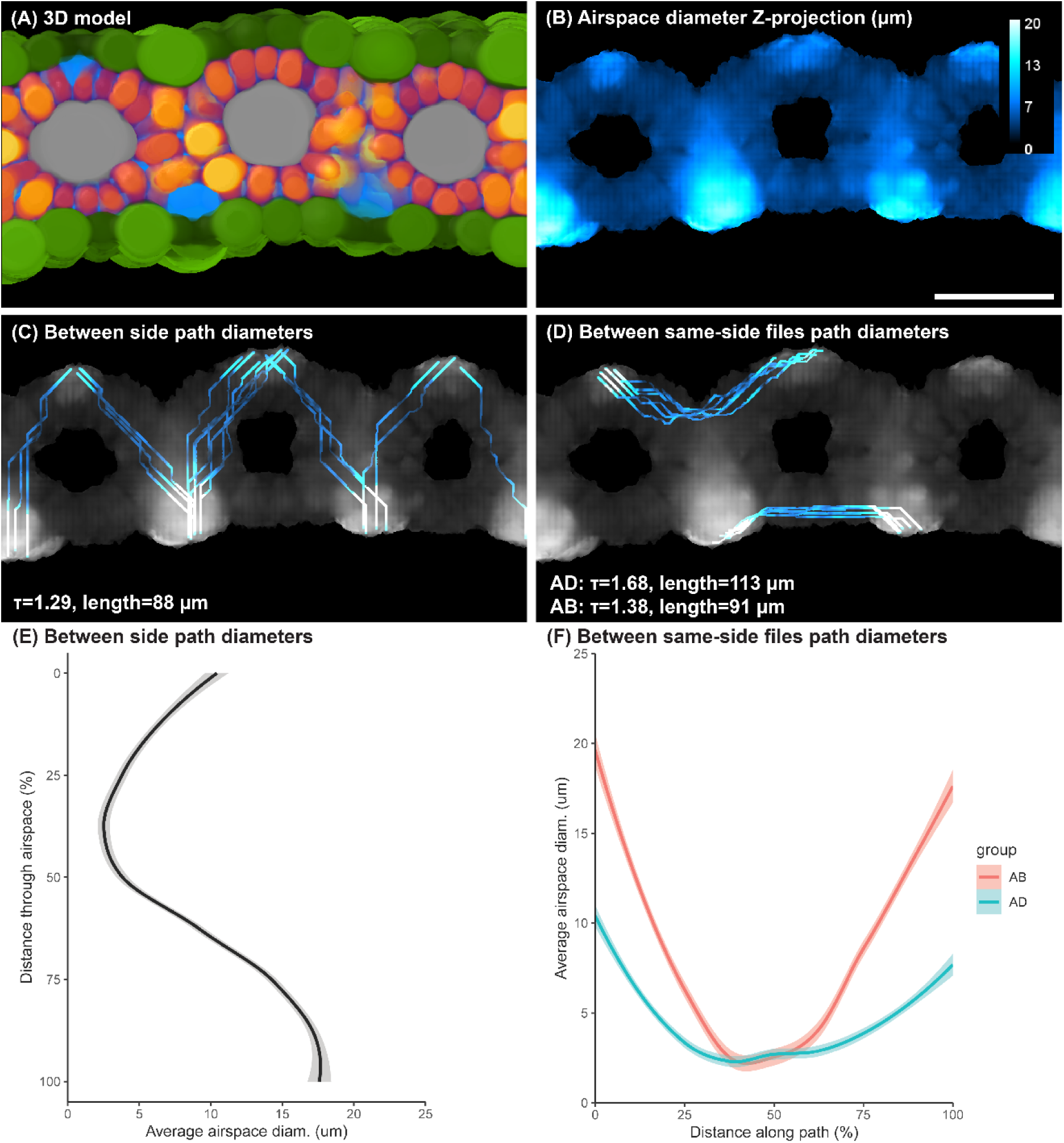
(A) 3D model of sub-sample shown in B-F (B) Z-projected airspace diameter of sub-sample (C) Shortest pathways from adaxial to abaxial stomata, color-coded to airspace diameter along pathway from B (D) Shortest pathways between same-side stomatal files, color-coded to airspace diameter along pathways from B (E) Plotted average diameter data across leaf depth from pathways in C (F) Plotted average diameter data along pathways in D.

**Supp. video 1:** Animation of wildtype sub-sample, showing differences in adaxial and abaxial morphology. Top animation is the adaxial side of the leaf, bottom animation is the abaxial side, and the middle animation is the cross-section of the sample. The epidermis animations are matched with the cross-sectional animation, so the user can scroll through the video and match the exact location of the scrolling epidermis edge with the shown cross-section scroll in real-time. Green LUT represents epidermis, orange LUT represents mesophyll, blue LUT represents airspace, and gray LUT represents vasculature. See Methods and Materials for explanation of machine-learning labeling and diameter labeling of tissue types.

